# Regulation of Parent-of-Origin Allelic Expression in *Arabidopsis thaliana* endosperm

**DOI:** 10.1101/521583

**Authors:** Karina S. Hornslien, Jason R. Miller, Paul E. Grini

## Abstract

Genomic imprinting is an epigenetic phenomenon set in the gametes prior to fertilization that causes differential expression of parental alleles mainly in the endosperm of flowering plants. The overlap between previously identified panels of imprinted genes is limited. In order to achieve high resolution sequencing data we have used sequence capture technology to investigate imprinting. Here, we present a bioinformatics pipeline to assay parent-of-origin allele specific expression and report more than 300 loci with parental expression bias. We find that the level of expression from maternal and paternal alleles in most cases is not binary, instead favouring a differential dosage hypothesis for the evolution of imprinting in plants. To address imprinting regulation, we systematically employed mutations in regulative epigenetic pathways suggested to be major players in the process. We establish the mechanistic mode of imprinting for more than 50 loci regulated by DNA methylation and Polycomb-dependent histone methylation. However, the imprinting patterns of the majority of genes were not affected by these mechanisms. To this end we also demonstrate that the RNA-directed DNA methylation pathway alone does not influence imprinting patterns in a substantial manner, suggesting more complex epigenetic regulation pathways for the majority of identified imprinted genes.

**Author summary:** Expression of gene copies only from the mother or the father’s genome, also termed imprinting, is a specialized epigenetic phenomenon that is found to be enriched at some genes expressed in the mammalian placenta and in the endosperm of the plant seed. Although several studies have reported on imprinted genes in plants, the identified loci are at large non-overlapping between reports. This motivated us to investigate in detail the expression pattern of imprinted genes in the endosperm and to determine how imprinting patterns are established at various imprinted loci. Although several underlying epigenetic regulation mechanisms have been demonstrated to establish imprinting patterns at certain genes, the majority of imprinted genes have not been linked to such mechanisms. In the present study we systematically investigated the mechanisms that are involved in establishing imprinting, by employing mutants of epigenetic regulators and high-throughput sequencing. In our high resolution study, we report more than 300 imprinted genes and demonstrate that the biological phenomenon imprinting involves gradual expression of parental gene copies rather than switching gene copies on or off. Notably, for the majority of imprinted genes, the mechanisms previously believed to be major to establish their imprinting patterns, are not responsible for mediating imprinting.

## Introduction

Fertilization in plants generates the seed, a structure with two fertilization products, the embryo, carrying the genetic makeup of the next generation, and the endosperm, a nourishing tissue that surrounds the embryo (Berger et al., 2006). The seed therefore consists of three genetically distinct components: the diploid embryo with one genome copy from each of the parents, the triploid endosperm with two maternal and one paternal copy, and a seed coat having the same genotype as the diploid mother-plant. The importance of balanced parental gene expression for proper development of progenies has been demonstrated in both mammals and plants (Barton et al., 1984; Birchler, 1993). In this respect, and in analogy to the mammalian placenta, the endosperm is a site of genomic imprinting; an epigenetic phenomenon that leads to parent-of-origin dependent expression of genes due to non-DNA sequence-based mechanisms established in the male and female germline (Feil and Berger, 2007 Nowack et al., 2010).

In mammals, imprinted genes are often involved in growth control (Leighton et al., 1995). In plants, the endosperm is the major tissue regulating the flow of nutrients to the embryo and is therefore a likely site for such effects. The evidence for similar function as in mammals has, however, been limited and it has been hypothesized that imprinting in plants may be a side effect of global epigenetic regulation taking place in the endosperm (Berger et al., 2012). Among widely used hypotheses to explain the selection of imprinted genes, one is the parent-conflict theory (Haig and Westoby, 2015) where allocation of resources is the major driver for evolution of parent-of-origin expression. Another is the differential dosage hypothesis (Birchler and Veitia, 2007) advocating that imprinting optimizes the abundance and expression of a transcript. A general discrepancy between these hypotheses is their expectation towards the types of genes that are imprinted and to what extent an allele is silenced (Rodrigues and Zilberman, 2015; Dilkes and Comai, 2004).

Several transcriptomics studies to identify imprinted genes, mostly focused on dicots such as *Arabidopsis* and cereals such as maize and rice, have resulted in lists ranging from 100 - 300 genes in different species (Gehring et al., 2011; Waters et al., 2011; Hsieh et al., 2011; Shirzadi et al., 2011; Luo et al., 2011; Wolff et al., 2011; Pignatta et al., 2014). Mutational analysis of these genes in many cases did not report any obvious seed phenotype (Masiero et al., 2011; Wolff et al., 2015; Shirzadi et al., 2011). The overlap of imprinted genes between different species and also between different experiments using the same species has been limited, suggesting methodical bias and stimulated speculation to whether imprinting undergoes rapid evolution (Gehring and Satyaki, 2017). Recent reports have enlarged the list of imprinted genes in the endosperm with a clear function (Costa et al., 2012; Chaudhury et al., 1997; Grossniklaus et al., 1998; Figueiredo et al., 2015). Analyzes utilizing higher tissue specificity and sequencing depth also report more evolutionary conservation of imprinted genes in close relatives (Hatorangan et al., 2016; Klosinska et al., 2016; Schon and Nodine, 2017; Moreno-Romero et al., 2017). Due to the limited overlap between large-scale screens for imprinted genes, however, the number of imprinted genes and the fraction of maternally and paternally biased genes is still controversial (Schon and Nodine, 2017).

Parent-of-origin-dependent expression is independent of the underlying DNA sequence and relies on epigenetic mechanisms which include DNA methylation and modifications of histones (Satyaki and Gehring, 2017; Raissig et al., 2011; Jullien et al., 2006). The Polycomb Repressive Complex 2 (PRC2) silences gene activity via methylation of lysine 27 on histone H3 (H3K27me3). In the *Arabidopsis thaliana* endosperm, PRC2 includes the histone methyltransferase (MTase) MEDEA (MEA) and the Su(Z)-homolog FERTILIZATION INDEPENDENT SEED 2 (FIS2), both of which show maternally biased expression (Thorstensen et al., 2011; Grossniklaus et al., 1998; Chaudhury et al., 1997). The corresponding genes were identified in screens for autonomous endosperm development, indicating that this complex also controls the gametophytic to sporophytic phase transition (Berger et al., 2006). Another important imprinting mechanism is DNA methylation, resulting from the activity of several DNA MTases, where METHYLTRANSFERASE 1 (MET1), the major *Arabidopsis* maintenance DNA MTase, sustains CG-methylation of hemi-methylated DNA after replication (Gehring et al., 2009). Removal of DNA methylation can be achieved either passively in the case of absence of MTase activity during DNA replication, or by an active mechanism involving DNA glycosylases such as DEMETER (DME) (Gehring, 2013; Penterman et al., 2007). In a simplistic model of genomic imprinting in plants, maternally expressed genes (MEGs) require the activity of DME on the maternal allele, whereas the paternal allele is repressed in the endosperm by MET1 activity prior to fertilization. Paternally expressed genes (PEGs) rely on PRC2 mediated H3K27me3 deposition to silence the corresponding maternal allele (Jullien and Berger, 2009; Köhler et al., 2004). Crosstalk between these epigenetic mechanisms has been reported. For instance, DNA methylation is hypothesized to block PRC2 histone methylation activity on paternal alleles whereas maternal alleles are repressed by PRC2, leading to expression in the presence of DNA methylation of the paternal allele in the case of certain PEGs (Weinhofer et al., 2010; Satyaki and Gehring, 2017). These generalizations are mainly based on correlations that have been drawn from genome-wide transcriptome and chromatin profiling studies. The mechanisms leading to imprinting have not been functionally tested for the majority of the reported imprinted genes.

In both animals and in plants, DNA methylation is found most frequently in the CG context. However, in plants, there is also extensive cytosine methylation in the CHG and CHH context (where H denotes all bases except G). *De novo* DNA methylation in all sequence contexts is established by the so-called RNA-directed DNA methylation (RdDM) (Wassenegger et al., 1994). The hallmark of RdDM activity is mirrored by methylation in the asymmetric CHH context and brought about by the MTase DOMAINS REARRANGED METHYLTRANSFERASE 2 (DRM2). CG, CHG and CHH methylation marks are maintained by MET1, CHROMOMETHYLASE 3 (CMT3) and CMT2, respectively (Zhang et al., 2013; Henderson and Jacobsen, 2007; Stroud et al., 2014). The canonical RdDM machinery in plants involves RNA polymerase IV (PolIV) recruitment to DNA (Law et al., 2013) and PolIV 25-30nt single stranded RNA molecules further transcribed to double stranded RNA by RNA DEPENDENT RNA POLYMERASE 2 (RDR2) (Zhai et al., 2015; Li et al., 2015). DICER LIKE (DCL) 3 cleaves the dsRNA to 24nt small interfering (si) RNA fragments that are loaded to ARGONAUTE 4 (AGO4) of the RNA-induced silencing complex (RISC) and guided to a newly produced RNA polymerase V (PolV) transcript where it associates with the MTase DRM2 (Cao and Jacobsen, 2002; Feng et al., 2010; Law and Jacobsen, 2010; Grimanelli and Roudier, 2013; Bond and Finnegan, 2007; Matzke et al., 2015). Traditionally, transcripts from RNA polymerase II (PolII) that generate small RNAs have been thought to act mainly in post transcriptional gene silencing (PTGS) as 21nt miRNA that interact with mRNA to infer degradation or translational inhibition (Lee et al., 2004; Ramachandran and Chen, 2008). Recent evidence, however suggests non-canonical forms of RdDM where single stranded RNA transcripts from PolII are made double stranded by RNA DEPENDENT RNA POLYMERASE 6 (RDR6) and further cleaved by DCL2 or DCL4 (Nuthikattu et al., 2013), loaded to AGO6 and guided to a PolV target site analogous to the canonical RdDM pathway. In a postulated model, RDR6-RdDM act as an initial silencer of active transposons, in a second step requiring PolIV-RdDM to maintain inactive transposons silent (Bond and Baulcombe, 2015; Panda et al., 2016).

Several lines of evidence suggest a role for small RNA and RdDM in imprinting. Small RNAs that map to imprinted genes have been identified both in *A. thaliana* and cereals (Rodrigues et al., 2013; Pignatta et al., 2014). For two maternally expressed loci, *SUPPRESSOR OF drm1 drm2 cmt3* (*SDC*) and *MOP9.5* (AT5G24240), PolIV activity have been demonstrated to be required for silencing of the paternal allele in the endosperm (Vu et al., 2013). Other components of the RdDM pathway, such as DCL4 have also been shown to relieve imprinting for a specific locus if mutated (Bratzel et al., 2012). The main subunit of PolIV is absent in the central cell, although CHH methylation can be detected (Vu et al., 2013; Ingouff et al., 2017), thus indicating a role for a non-canonical RdDM pathway (Satyaki and Gehring, 2017). Recent evidence suggests a regulatory role for RdDM in the fertilized seed, particularly in endosperm tissues. A domain in the endosperm termed peripheral endosperm displays decondensed chromatin and typically lack of chromocenters (Baroux et al., 2007). In pollen, similar chromatin decondensation has been coupled to increased RdDM activity (Schoft et al., 2009) and the abundant presence of small RNAs in seeds have therefore been suggested to stem from decondensed endosperm chromatin (Mosher et al., 2009).

Previously reported panels of imprinted genes show limited overlap (Wolff et al., 2011; Shirzadi et al., 2011; Gehring et al., 2011; Hsieh et al., 2011). In a comparison of three datasets based on RNASeq SNP detection, only three common genes were identified as imprinted. The addition of a fourth seed transcriptome based on the functional absence of a paternal genome in the endosperm (Shirzadi et al., 2011; Nowack et al., 2006; Aw et al., 2010) did not overlap with genes classified as imprinted. In this study, we selected a set of 1011 genes that had previously been shown to be imprinted (Wolff et al., 2011; Gehring et al., 2011; Hsieh et al., 2011), genes considered regulators of imprinting/control genes, as well as a set of genes regulated in the absence of paternal contribution to the endosperm (Shirzadi et al., 2011). We developed a bioinformatics process that we call the Informative Reads Pipeline (IRP) to analyze RNAseq data and detect parent-of-origin allele specific expression. Using three different ecotypes of *A. thaliana*, we demonstrate that the IRP identifies parent-of-origin dependent expression preference. We report more than 300 genes with maternal or paternal bias in reciprocal crosses in at least two ecotypes. We find that the level of expression from maternal and paternal alleles of imprinted genes to be relative rather than absolute, thus favouring the differential dosage hypothesis for the evolution of imprinting. To address the regulation of these imprinted genes, we have analyzed imprinted expression in crosses involving mutants in the MET1, PRC2 and RdDM pathways. We report that even though the majority of paternally expressed genes are imprinted by PRC2, only one third of the maternally biased genes can be explained by MET1, thus suggesting the involvement of further mechanisms. To this end we have investigated the role of RdDM, and found that only a few of the genes tested were significantly affected by the absence of components of RdDM. Our results suggest more complex epigenetic regulation pathways for the majority of identified imprinted genes.

## Results

We performed transcriptome analysis of parent-of-origin allele specific expression in reciprocal crosses between accessions that allow allelic detection due to prominent SNP variation (Ossowski et al., 2008; Schneeberger et al., 2011). Here we employ three different accessions of *A. thaliana*; Columbia-0 (Col-0), Landsberg *erecta* (L*er*) and Tsushima (Tsu) in reciprocal crosses. To verify that Col-0/L*er* and Col-0/Tsu-1 self and reciprocal crosses were comparable, seeds from crosses were inspected four days after pollination (4DAP, Figure 1A). Although a slight variation in seed size and developmental stage could be observed, thorough staging showed that in all crosses most seeds (>65%) had uniformly reached the early globular stage (Figure 1B, STable 1). Total RNA from seeds were harvested from dissected siliques at 4DAP and reverse transcribed to cDNA. In order to achieve high read depth and also to assess low abundance transcripts, Nimblegen custom made probes covering all exons of selected target genes were utilized to extract targets for sequencing (see Materials and Methods). To take full advantage of the sequence capture technology, target amplicons were reduced from a full transcriptomic study to a selected group of target amplicons. The 1011 targets selected for probe capture were genes that had previously been shown to be imprinted (Wolff et al., 2011; Gehring et al., 2011; Hsieh et al., 2011), genes considered regulators of imprinting/control genes, as well as a set of genes regulated in the absence of paternal contribution to the endosperm (Shirzadi et al., 2011). The target set cover 78% of all published imprinted genes from whole genome studies (Wolff et al., 2011; Gehring et al., 2011; Hsieh et al., 2011; Pignatta et al., 2014) The probe capture method provided high resolution sequencing data and generated 1.06x10^12^ bases in over 3.53x10^9^ paired end reads. The number of pairs per sample ranged from 24.6x10^6^ to 42.1x10^6^, with 32.7x10^6^ as the average. The number of pairs per cross ranged from 151x10^6^ to 244x10^6^, with 196x10^6^ as the average (see SData 2)

**Figure 1:**
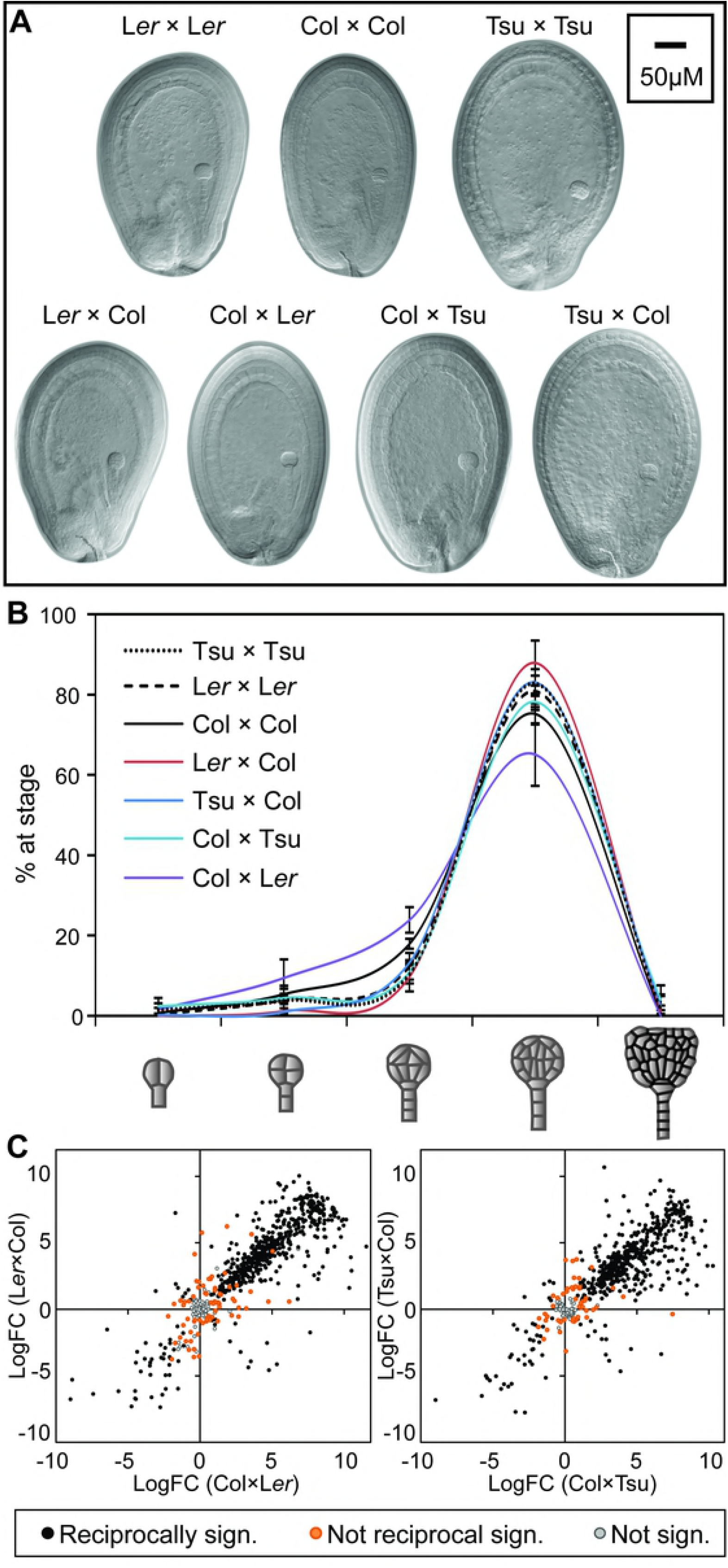
Imprinting in *Arabidopsis* ecotypes Col-0, Tsu-1 and L*er* at four days after pollination. **A)** Seeds from crosses within and between ecotypes used in this study were investigated at four days after pollination (DAP) indicating that there is no obvious difference regarding seed development depending on the specific ecotype used as a maternal or paternal contributor in a cross. **B**) In >65% of all seeds investigated, the embryos were classified as 32-cell to globular stage at 4DAP, and only very few deviated from this stage. Data is presented as means ±standard deviation across three biological replicates. **C)** cDNA libraries from RNA harvested from 4DAP tissue were sequenced, and informative reads were used to detect significant (sign.) parentally biased expression from two ecotype cross pairs; Col-L*er* and Col-Tsu. Cross direction is always indicated as ♀ x ♂.

For the purpose of detection of allele specific expression, we developed the Informative Reads Pipeline (IRP) (SFigure 1). In contrast to conventional SNP analysis, i.e. weighing reads that map to published SNP libraries, we have utilized all SNP and InDel variation that we observed empirically in all transcripts. Read pairs from the three homozygous wild type crosses (Col-0 x Col-0, L*er* x L*er* and Tsu x Tsu) were used to generate a strain-specific consensus sequence for each of the 1011 genes in our study. Read pairs from all crosses were mapped to our new consensus sequences (SData 2, see Materials and Methods for details). Reads from Col-0 x Col-0 and L*er* x L*er* crosses were mapped to the Col and L*er* consensus sequences, with each read allowed to map to up to 2 target sequences. This was repeated similarly for reads from Col-0 x Col-0 and Tsu x Tsu crosses. A pair was considered “informative” if it mapped to both strain versions of the same gene, and only one mapping was InDel-free, or both mappings were InDel-free but one contained fewer mismatches (i.e. SNPs). Overall, 640 million (8.6%) reads became informative reads, covering 776 (76,7%) of our 1011 targets.

Homozygous crosses were used to filter genes for which allelic expression would not be detectable by informative reads (i.e. reads covering ecotype specific SNP and/or InDels). Genes were filtered if they provided insufficient informative reads counts or if their informative reads did not sufficiently favor the true parent (SFigure 2). In total, 718 genes were retained for the Col vs. L*er* comparisons and 627 for the Col vs. Tsu comparisons. There were 550 genes in the intersection of these two sets (see SData 3).

Informative reads from heterozygous crosses were used to detect differential expression between the parental alleles (see Materials and Methods for details). Since the endosperm contains a 2:1 ratio of maternal to paternal genomes, maternal reads were halved to ease the detection of deviation to a 1:1 ratio through the differential expression analysis. Genes were designated as biased if they showed significant excess (p<0.05) in maternal or paternal expression, respectively. In L*er* x Col and Col x L*er* crosses, 590 and 582 genes, respectively, showed significant parental preference. In Tsu x Col and Col x Tsu crosses 525 and 521 genes, respectively showed significant parental preference (Figure 1C). 549 and 497 genes were significantly parentally biased in Col-L*er* and Col-Tsu reciprocal crosses (Figure 1C).

In order to compare IRP to conventional SNP analysis, reads mapping to a single SNP was chosen for each gene and differential expression between maternally and paternally originating reads was calculated. The obtained SNP fold change (FC) was compared to the IRP FC for the same genes in the same samples (SFigure 3). The analysis show high correlation, with a correlation coefficient close to 1 for all crosses, thus verifying that the accuracy of IRP correlates to traditional SNP mapping. Some outliers were observed, e.g. FIS2 had opposing FC in SNP vs IRP in Col-Tsu crosses. Meticulously analyzing reads mapping to the selected SNP and other SNPs in the same gene show that the selected SNP attracted far more reads than other SNPs within the same gene, suggesting that during library preparation some regions may have been amplified to greater extent than others. This is unlikely due to identical reads, originating from other loci that are mapping to this region, as we used a sequence capture approach with unique probes (SData 4). By IRP, the mapping of such reads can only be partially avoided, but by using all informative reads across a transcript (all SNPs and InDels) this effect is largely diminished. In this respect, the IRP has an advantage since in our setup, InDels comprise 26,5% and 34,0% of the Col-L*er* and Col-Tsu variation in targets with informative reads, respectively, the remaining variation is due to SNPs. A limited number of genes have only InDels and no SNPs; 1,7% (14 genes) in Col-L*er* and 4,5% (35 genes) in Col-Tsu.

RNA samples extracted from whole seeds can lead to biased maternal to paternal ratio as it contains the maternally derived seed coat. Genes can be incorrectly identified as maternally expressed imprinted genes when contributed from maternally derived tissues or transcripts of pre-fertilization, maternal gametophytic origin. In a recent report, it has been argued that only confined preferential expression in the endosperm allows for identification of truly imprinted maternally expressed genes (Schon and Nodine, 2017). In order to correct for maternal bias, we have generated conservative and moderate stringency groups for further analysis (See SMethods). The conservative group harbours targets that show restricted expression pattern to the endosperm, while in the moderate and low stringency groups expression in other tissues is tolerated as long as it does not exceed endosperm expression substantially. In all groups known seed coat specific genes where filtered away.

After filtering (described above and Materials and Methods) a conservative group was defined consisting of genes with confined preferential expression to the endosperm as previously described (Schon and Nodine, 2017). Only 85 and 84 transcripts from Col-L*er* and Col-Tsu, respectively, were amongst our targets with sufficient informative read information, and are the basis of our conservative endosperm specific group. A moderate stringency group was on the other hand defined based on previously reported seed coat enrichment (Schon and Nodine, 2017; Belmonte et al. 2011). Here, we repeated the test for enrichment in seed tissues especially emphasised on removing seed coat expressed genes while still accepting that a gene may be expressed in several tissues and still be imprinted in endosperm (SData 5, SMethods). Following these considerations, a moderate stringency set having support in previous studies not to be significantly enriched in the seed coat was generated consisting of 429 and 396 genes (including also the conservative endosperm enriched genes) in Col-L*er* and Col-Tsu respectively (SData 6).

A low stringency group was also defined for our target genes (SMethods). This group also included genes that do not have any expression profile available (due to missing coverage on the ATH1 microarray chip), arguing that we do not need prior knowledge on a gene’s expression pattern to identify paternally expressed imprinted genes. Paternally biased genes are unlikely a consequence of seed coat contamination and can still be selected for the analysis of their imprinting status. Distribution of FC in all targets and low stringency targets versus moderate targets show that genes filtered away indeed affect the FC values of MEGs and are a potential source of creating false positives when detecting MEGs from whole seed assays (SFigure 4).

### Detection of Loci with Parental Expression Bias

In the conservative endosperm specific group only 13 and 16 genes were identified as having reciprocally maternally biased expression in the Col-L*er* and Col-Tsu pair, respectively (p<0,01).Eight and nine genes were identified as having reciprocally paternally biased expression in the Col-L*er* and Col-Tsu pair (SFigure 5A, SData 7). 39 genes were reciprocally defined as to not have any parental bias in either direction for Col and L*er*, and 38 genes for Col and Tsu (SFigure 5A). Some genes have only parental bias in one direction of the cross and some are detected as having opposite parental bias patterns depending on the direction of the cross.

In the low stringency group there is a drastic increase in the number of maternally biased genes (SFigure 5B), reflecting a putative higher occurrence of false positives in this group as discussed previously. More than 300 genes were significantly assigned as having maternally biased expression in all crosses (p<0,01). Paternally biased expression was limited to 38 and 35 reciprocal genes for the Col-L*er* and Col-Tsu cross pairs respectively (p<0,01, SData 7).

In the moderate stringency group, we identified 265 genes that were preferentially maternally expressed in both Col-L*er* crosses and 32 genes that were preferentially paternally expressed in both crosses (p<0,01, Figure 2A, SData 7). In the Col-Tsu reciprocal crosses 253 and 31 genes were identified as being preferentially maternally or paternally expressed, respectively. 67 of the 429 genes in the Col-L*er* pair and 56 out of the 396 genes in the Col-Tsu pair were assigned as having no preferential parental bias (SData 7). Some genes showed only maternal or paternal bias in one cross direction while other genes had opposite preferential bias in the genes analyzed. The genes that have opposite preferential parentally biased expression may be a result of different level of expression in the ecotypes used as parents. E.g. a Col-allele may be stronger expressed than a L*er*-allele independent of its parental origin. To this end, the differential expression analysis was repeated to incorporate read counts from ERCC spike-in sequences. This analysis was performed with the goal of indicating whether absolute expression level differences between samples had induced false-positive or false-negative expression differences (see Materials and Methods for details). More than half of opposite preferentially biased genes (e.g. MEG in Col x L*er* but PEG in L*er* x Col) were found to be more than three-fold differentially expressed in Col versus L*er* or Tsu (STable 2, SData 8). The group that only show parental bias in one direction may be candidates for an ecotype specific parental expression pattern and in our setup of ecotypes we were able to verify that some of these genes do indeed behave similarly in the different crosses depending on the ecotype used as maternal or paternal contributor (STable 3).

**Figure 2:**
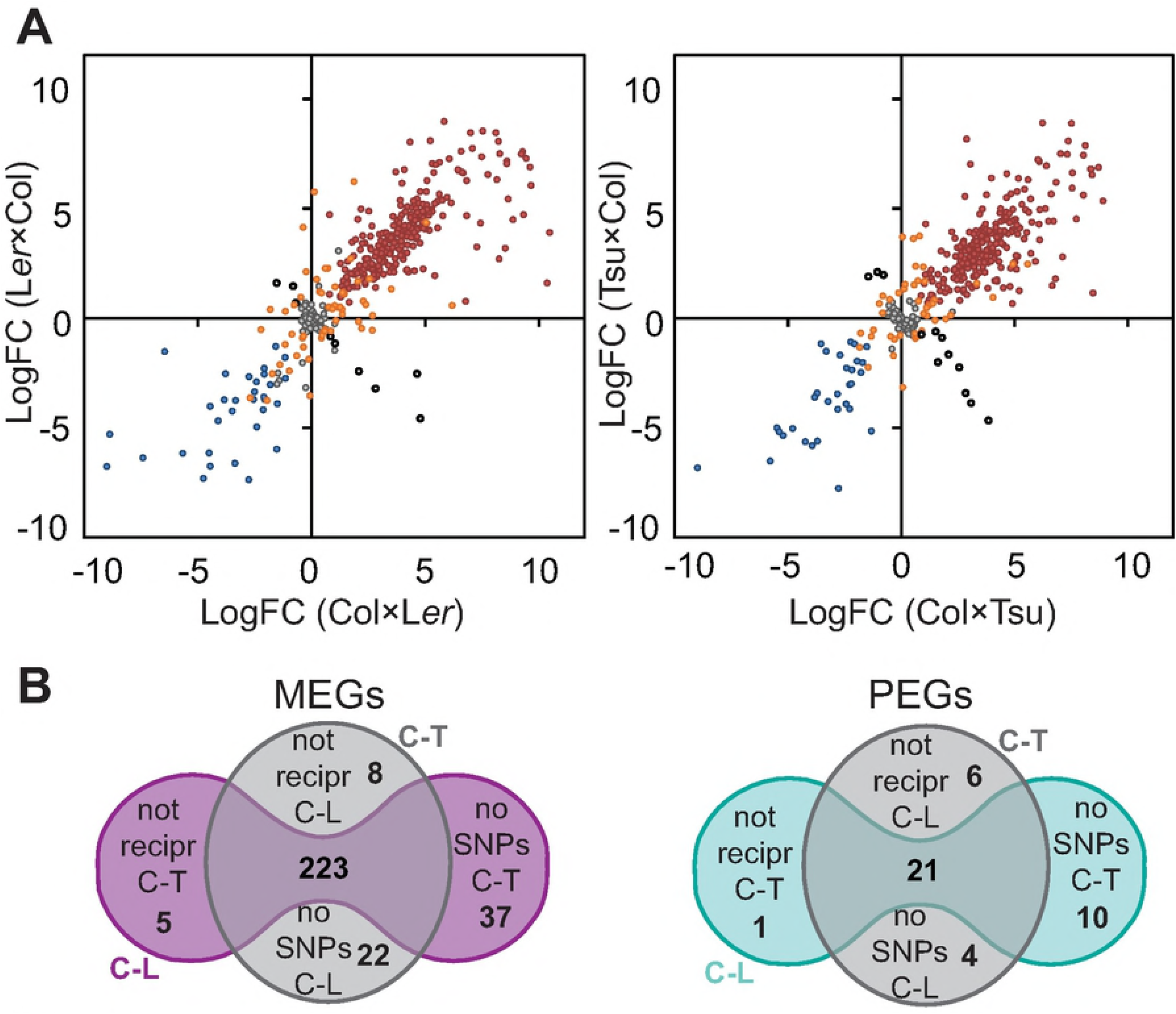
Parentally biased expression in the moderate stringency group. **A)** The LogFC between maternal and paternal reads for each gene in each cross was plotted against its reciprocal counterpart to display preferential mapping, i.e. parentally biased expression. A positive LogFC signifies maternal preference while a negative LogFC signifies paternal preference. The moderate stringency group of targets for the two cross pairs; Col-L*er* (left panel) and Col-Tsu (right panel). Grey filled circles signify targets that do not have significant parentally biased expression. Red and blue filled circles have maternally or paternally biased expression, respectively, in both directions of a cross pair. Orange filled circles have parentally biased expression in only one direction of a cross pair. Black circles show opposite preferential expression depending on the direction of the cross. Cross direction is always indicated as ♀ x ♂. **B)** Great overlap is observed between preferentially expressed genes detected in all ecotype crosses used in this study; Col-L*er* (C-L) reciprocal crosses and Col-Tsu (C-T) reciprocal crosses, Maternally Expressed Genes (MEGs) in left panels and Paternally Expressed Genes (PEGs) in right panels. 244 genes show consistent preferential bias in our moderate stringency group. Genes that do not overlap are most frequently genes that do not have sufficient SNP/InDels in the alternative cross pair and could not be assessed. Few genes show inconsistent or not reciprocal preferential expression in the two cross pairs used in this study. Grey circle; genes that show reciprocal parent preferential expression in Col-Tsu cross pair. Coloured peanut; genes that show reciprocal parent parental bias in Col-L*er* cross pair.

### Most Parentally Biased Genes Are Conserved in Ecotypes

Although some of the identified genes with parental preferential expression do not have SNP coverage in all ectypes, we do observe a high degree of overlap between the ecotypes. Very few genes that have SNPs/InDels in all ecotypes show inconsistent or not reciprocal preferential expression in the alternate cross pair. In the conservative endosperm-specific group only 7 genes (28%) did not show reciprocal imprinting in both cross pairs. Only 3 genes in the conservative group were not possible to evaluate in the alternate cross pair due to lack of SNP/InDel coverage or insufficient read counts to pass the initial filtering step. 18 genes (72%) showed consistent parental preferential expression (SFigure 5C, SData 7). In the moderate stringency group only a total of 20 genes (7,5 %) do not have the same preferential expression pattern reciprocally. A total of 73 genes (28%) do not have SNP/InDel coverage in the alternate cross pair and can therefore not be evaluated in both Col-L*er* and Col-Tsu pairs, while 244 genes (92%) overlap between all ecotypes. Due to the high consistency between ecotypes we regarded it as “safe” to include these 73 in further analysis since they were reciprocally imprinted in at least one cross pair (Figure 2B, SData 7). A similar trend was observed by analysis of the low stringency dataset. Of the genes that have SNP/InDel coverage in both alternate cross pairs, 92% (of 322) have the same preferential expression pattern reciprocally (SFigure 5D, SData 7). In this set of genes, bias due to seed coat expression could not be excluded, and it was mainly included to screen for transcripts that were paternally biased and to measure the number of false negatives compared to the moderate stringency group. Indeed, we observed a higher number of candidate maternally biased genes, whereas the number of paternally biased genes only increased by two genes. The maternal dataset was not further considered due to lack of seed coat expression data, but the identified paternally biased genes were added to the list of paternally biased genes but not included in further analysis (Table 1).

**Table 1:** Imprinted Paternally Expressed Genes (PEGs) detected in the moderate stringency group.

| AGI | Annotation | Short Description | H | G | W | S | P | ColxLer<br>FC | LerxCol<br>FC | ColxTsu<br>FC | TsuxCol<br>FC | Expression<br>pattern | # ecotypes |
| --- | --- | --- | --- | --- | --- | --- | --- | --- | --- | --- | --- | --- | --- |
| AT1G66630 | AT1G66630 | Protein with RING/U-box |  |  | x |  |  | -6,82 | -164,92 | -6,85 | -216,13 | ess. >8-fold | CLT |
| AT3G50720 | AT3G50720 | Protein kinase superfamily protein |  |  | x | d |  | -513,55 | -108,53 | -507,23 | -112,84 | ess. >8-fold | CLT |
| AT1G60400 | AT1G60400 | F-box/RNI-like superfamily |  |  | x | d |  | -22,04 | -107,27 | -18,78 | -49,33 | ess. >8-fold | CLT |
| AT1G34650 | HDG10 | Homeodomain GLABROUS 10 |  |  | x | d |  | -22,07 | -16,11 | -13,00 | -48,49 | ess. >8-fold | CLT |
| AT1G65360 | AGL23 | AGAMOUS-like 23 |  |  | x |  |  | -17,30 | -25,67 | -37,15 | -40,84 | ess. >8-fold | CLT |
| AT5G54350 | AT5G54350 | C2H2-type zinc finger protein |  |  | x |  |  | -50,77 | -71,13 | -41,90 | -36,00 | ess. >8-fold | CLT |
| AT2G20160 | MEO | E3 ubiquitin ligase SCF complex subunit |  |  | x |  |  | -5,40 | -7,49 | -3,43 | -5,53 | ess. >8-fold | CLT |
| AT4G10160 | AT4G10160 | RING/U-box superfamily protein |  |  | x |  |  | -13,78 | -5,76 | -9,93 | -2,83 | ess. >7-fold | CLT |
| AT1G67820 | AT1G67820 | Protein phosphatase 2C family |  | x |  |  | x | -14,32 | -13,18 | -13,03 | -10,70 | ess. >6-fold | CLT |
| AT1G49290 | AT1G49290 | Rho guanine nucleotide exchange factor |  |  | x |  |  | -27,08 | -157,69 | -54,83 | -90,79 | ess. >5-fold | CLT |
| AT3G49770 | AT3G49770 | Hypothetical protein |  |  | x |  | x | -172,24 | -82,64 | -44,79 | -31,81 | ess. >5-fold | CLT |
| AT1G48910 | YUC10 | Flavin-containing monooxygenase | x | x | x |  | x | -10,22 | -13,41 | -5,39 | -15,13 | ess. >5-fold | CLT |
| AT2G36560 | AT2G36560 | AT hook motif DNA-binding | x | x |  |  |  | -5,30 | -31,05 |  |  | ess. >4-fold | CL |
| AT4G23110 | AT4G23110 | Insulin-like growth factor |  | x |  |  |  | -10,37 | -98,22 |  |  | ess. >4 fold | CL |
| AT5G50470 | NF-YC7 | Nuclear factor Y, subunit C7 |  |  | x |  |  | -22,75 | -70,35 |  |  | ess. >3 fold | CL |
| AT5G26700 | AT5G26700 | RmlC-like cupins superfamily protein |  | M |  |  |  | -4,23 | -5,82 |  |  | ess. >1-fold | CL |
| AT1G65330 | PHE1 | MADS-box transcription factor |  |  | x | d |  |  |  | -7,04 | -10,97 | ess. >8-fold | CT |
| AT3G19160 | IPT8 | ATP/ADP isopentenyltransferase |  |  | x |  |  |  |  | -6,52 | -3,20 | ess. >7-fold | CT |
| *AT3G11310 | AT3G11310 | Myb/SANT-like DNA-binding |  | x |  |  | x | -5,24 | -63,51 | -21,67 | -63,22 | not on ATH1 | CLT |
| *AT5G54360 | AT5G54360 | C2H2-like zinc finger protein |  |  |  | d |  | -22,92 | -103,61 | -10,83 | -210,99 | not on ATH1 | CLT |

Fold Change (FC) values are given as calculated by differential expression (DE) of the maternal and paternal allele. P-value range obtained from DE analysis ranged from 1,15x10<sup>-7</sup> to 0,001157.
Imprinting state reported in selected whole genome surveys of imprinting (x if consistent with this data): H= Hsieh et al., 2011; G= Gehring et al., 2011; W= Wolff et al., 2011; P= Pignatta et al., 2014; S= Shirzadi et al., 2011 downregulated (d) in *cdka; l*-screen. M=reported as MEG.
Expression pattern indicated is based on the microarray ATH1-chip library (Belmonte et al., 2013) and re-analysis (see Material and Methods) for tissue enrichment to identify endosperm specific (ess.) expression compared to all other seed compartments at globular stage.
CLT= SNPs in all ecotypes tested, CL= only SNP between Col-0 and Ler, CT= Only SNP between Col-0 and Tsu.
\*Indicate PEGs included from the low stringency group.

### Genomic Imprinting Is a Quantitative Phenomenon

In the conservative dataset, we could verify a total of 18 genes that were parentally biased in reciprocal crosses in three ecotypes (STable 4). This included 11 MEGs and 7 PEGs counting previously well studied imprinted genes such as *MATERNALLY EXPRESSED PAB C-TERMINAL* (*MPC*), *HOMEODOMAIN GLABROUS 10* (*HDG10*) and *MEIDOS* (*MEO*). As observed in our dataset and also reported previously (Schon and Nodine, 2017), the number of genes with maternal expression bias is strongly reduced compared to other whole genome studies (Wolff et al., 2011; Gehring et al., 2011; Hsieh et al., 2011; Pignatta et al., 2014) However, there are no overlapping loci when comparing a torpedo stage set of imprinted genes reported by these authors (Schon and Nodine, 2017) and our conservative dataset at globular stage (SFigure 6), although they are characterized as imprinted in other studies (Tiwari et al., 2008; Wolff et al., 2011; Gehring et al., 2011; Hsieh et al., 2011).

Analysis of our moderate stringency dataset identified a total of 35 genes with preferential paternal expression (STable 5). All displayed preference for all biological replicates in reciprocal crosses between an ecotype pair and 21 could be detected between three ecotypes (Figure 2B, STable 5). Although the majority of these genes displayed more than a five-fold preferential expression from the paternal genome, we observed a wide range of parental expression ratios, ranging from two to several hundred (STable 5). We therefore conclude that genomic imprinting is a quantitative phenomenon and refer to genes that display significant differential expression from the parental genomes as imprinted. Most of the paternally expressed genes (PEGs, Table 1) have support as being endosperm specific compared to other seed compartments (Schon and Nodine, 2017) and from our re-analysis of the Goldberg-Harada dataset (Belmonte et al., 2011) (SData 5).

In conclusion, the low stringency group has a high potential of false positive MEG discovery, whereas the conservative endosperm enriched group has high potential of false negative MEGs and PEGs detection and exclude genes expressed in the seed coat from being discovered as imprinted genes. For further analysis of imprinting, we focussed on the moderate stringency group; this group contains the conservative endosperm enriched imprinted genes, and transcript enriched in equal amounts or significantly less in the seed coat than in the endosperm.

### Maternally Biased Genes Are Identified More Frequently

Analysing the same dataset for maternally expressed genes (MEGs) identified a total of 282 genes with a significant (p<0,01) maternal preference in all replicates and cross directions. 80% of these (223) were present in three ecotypes (Figure 2B, STable 6). Comparable to the PEG analysis, FC expression ratios ranged from small, but significant changes to several hundred-fold. A majority of the MEGs showed a maternal preference higher than 10-fold, for instance 142 genes have more than 10-fold maternal preference in both reciprocal crosses of the L*er*-Col ecotype pair. Even in a worst-case scenario, where seed coat expression constitutes 50% of the expression in the seed (ANOVA <1), a MEG FC higher than 5 would be significant.

Taken together we could verify imprinting in the globular seed stage for almost half of previously reported genes selected in our setup (Hsieh et al., 2011; Gehring et al., 2011; Wolff et al., 2011) (Figure 3). Our dataset also include a set of deregulated genes in the absence of paternal contribution to the endosperm (Shirzadi et al., 2011) and genes involved in epigenetic regulation. From this set of genes, 260 genes could be tested in the moderate stringency group and 161 could be verified to be parentally biased (STable 6). This subset of genes also identified 119 genes with maternal preferential expression (p<0,01) not previously shown to be imprinted (STable 6). One third of these genes displayed a ten-fold maternal preference in all replicates in three ecotypes and was thus regarded as highly significant novel MEGs (Table 2).

**Figure 3:**
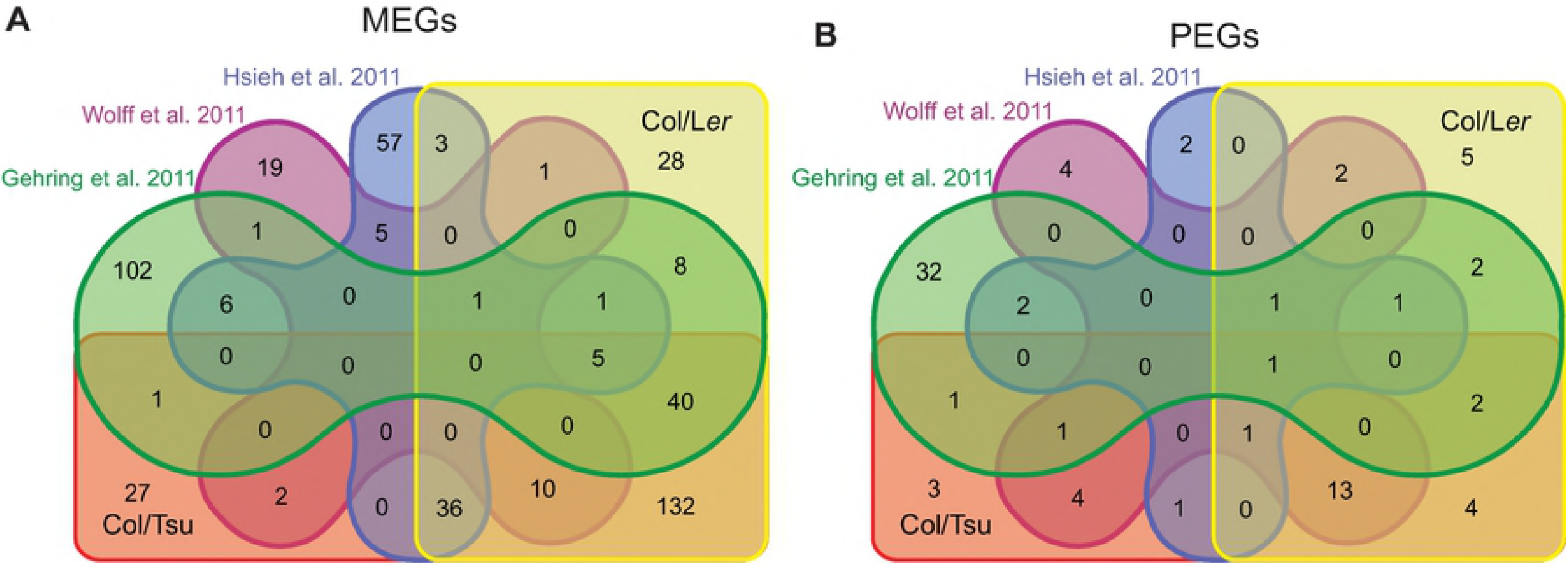
Parental biased expression is consistent with previous reported imprinting state. Intersection of parental biased expression reported in the moderate stringency group in this study and previous reports (Gehring et al., 2011; Hsieh et al., 2011; Wolff et al., 2011) for **A)** maternally expressed genes (MEGs) and **B)** paternally expressed genes (PEGs).

**Table 2:** Selected novel imprinted maternally express.ed genes (MEGs) detected in the moderate stringency set with Fold change (FC) > 10.

| AGI | Gene name | Short Description | H | G | W | S | P | ColxLer<br>FC | LerxCol<br>FC | ColxTsu<br>FC | TsuxCol<br>FC | Expression<br>pattern | # ecotypes |
| --- | --- | --- | --- | --- | --- | --- | --- | --- | --- | --- | --- | --- | --- |
| AT4G00730 | ANL2 | Homeobox-leucine zipper family protein |  |  |  | x |  | 22,80 | 15,20 | 21,35 | 10,78 | ess. >4-fold | CLT |
| AT4G32850 | nPAP | Nuclear poly(a) polymerase |  |  |  | x |  | 15,25 | 15,93 | 11,72 | 13,65 | ess. >4-fold | CLT |
| AT2G34200 | AT2G34200 | RING/FYVE/PHD zinc finger superfamily |  |  |  | x |  | 14,89 | 41,79 | 24,78 | 45,18 | ess. >3-fold | CLT |
| AT4G39390 | NST-K1 | Nucleotide sugar transporter-KT1 |  |  |  | x |  | 20,93 | 17,25 | 19,86 | 13,72 | ess. >3-fold | CLT |
| AT1G27000 | AT1G27000 | GRIP/coiled-coil protein, putative (DUF1664) |  |  |  | x |  | 14,24 | 14,62 | 19,33 | 13,86 | ess. >2-fold | CLT |
| AT4G00550 | DGD2 | Digalactosyl diacylglycerol deficient 2 |  |  |  | x |  | 21,79 | 17,34 | 12,50 | 18,12 | ess. >2-fold | CLT |
| AT1G22040 | AT1G22040 | Galactose oxidase/kelch repeat |  |  |  | x |  | 24,39 | 26,11 | 22,95 | 15,87 | ess. >3-fold<br>PG | CLT |
| AT2G32990 | GH9B8 | Glycosyl hydrolase 9B8 |  |  |  | x |  | 184,28 | 370,19 | 67,87 | 140,99 | ess. >7-fold<br>LC | CLT |
| AT1G19140 | AT1G19140 | Ubiquinone biosynthesis COQ9-like protein |  |  |  | x |  | 18,55 | 13,48 | 17,99 | 17,66 | ess. >2-fold<br>BC | CLT |
| AT1G17950 | MYB52 | Myb domain protein 52 |  |  |  | x |  | 802,46 | 66,50 | 323,47 | 111,52 | ND in seed | CLT |
| AT1G43800 | FTM1 | Plant stearyl-acyl-carrier-protein |  |  |  | x |  | 39,64 | 132,24 | 48,18 | 79,64 | ND in seed | CLT |
| AT1G69530 | EXPA1 | Expansin A1 |  |  |  | x |  | 599,29 | 180,23 | 182,17 | 474,81 | ND in seed | CLT |
| AT1G71940 | AT1G71940 | SNARE associated Golgi protein family |  |  |  | x |  | 18,66 | 13,70 | 25,17 | 14,70 | ND in seed | CLT |
| AT1G79530 | GAPCP-1 | Glyceraldehyde-3-phosphate dehydrogenase |  |  |  | x |  | 29,61 | 24,29 | 23,23 | 22,17 | ND in seed | CLT |
| AT2G37390 | NAKR2 | Chloroplast-targeted copper chaperone |  |  |  | x |  | 126,68 | 351,58 | 104,94 | 21,53 | ND in seed | CLT |
| AT2G46650 | CB5-C | Cytochrome B5 isoform C |  |  |  | x |  | 56,82 | 501,76 | 65,32 | 150,57 | ND in seed | CLT |
| AT3G07190 | AT3G07190 | B-cell receptor-associated protein 31-like |  |  |  | x |  | 23,67 | 17,73 | 22,66 | 10,21 | ND in seed | CLT |
| AT3G25540 | LAG1 | TRAM, LAG1 and CLN8 (TLC) lipid-sensing |  |  |  | x |  | 15,12 | 35,33 | 12,50 | 15,42 | ND in seed | CLT |
| AT3G53000 | PP2-A15 | Phloem protein 2-A15 |  |  |  | x |  | 24,97 | 13,21 | 11,95 | 11,68 | ND in seed | CLT |
| AT3G55320 | ABCB20 | P-glycoprotein 20 |  |  |  | x |  | 53,45 | 41,84 | 44,38 | 30,86 | ND in seed | CLT |
| AT4G00780 | AT4G00780 | TRAF-like family protein |  |  |  | x |  | 642,63 | 26,27 | 416,02 | 118,11 | ND in seed | CLT |
| AT4G12590 | AT4G12590 | ER membrane protein complex subunit-like |  |  |  | x |  | 15,39 | 15,52 | 13,08 | 12,92 | ND in seed | CLT |
| AT4G21760 | BGLU47 | Beta-glucosidase 47 |  |  |  | x |  | 781,39 | 118,27 | 187,41 | 99,14 | ND in seed | CLT |
| AT4G26640 | WRKY20 | WRKY family transcription factor |  |  |  | x |  | 26,62 | 26,72 | 36,82 | 17,19 | ND in seed | CLT |
| AT4G29530 | AT4G29530 | Pyridoxal phosphate phosphatase-related |  |  |  | x |  | 14,83 | 16,82 | 29,43 | 21,82 | ND in seed | CLT |
| AT4G31770 | DBR1 | Debranching enzyme 1 |  |  |  | x |  | 18,40 | 13,17 | 10,18 | 11,50 | ND in seed | CLT |
| AT5G28530 | FRS10 | FAR1-related sequence 10 |  |  |  | x |  | 20,43 | 26,67 | 18,59 | 12,89 | ND in seed | CLT |
| AT5G47790 | AT5G47790 | SMAD/FHA domain-containing protein |  |  |  | x |  | 30,26 | 17,17 | 18,13 | 10,54 | ND in seed | CLT |
| AT4G31080 | AT4G31080 | Integral membrane metal-binding (DUF2296) |  |  |  | x |  | 39,29 | 29,17 | 24,62 | 24,83 | NA | CLT |
| AT5G14620 | DRM2 | Domains rearranged methyltransferase 2 |  |  |  |  |  | 26,20 | 38,15 | 46,43 | 15,11 | NA | CLT |

Fold Change (FC) values are given as calculated by differential expression of the maternal and paternal allele. P-values obtained from DE analysis ranged from $2.57 \times 10^{-8}$ to 0,0030.
Imprinting state reported in selected whole genome surveys of imprinting: H= Hsieh et al., 2011; G= Gehring et al., 2011; W= Wolff et al., 2011; P= Pignatta et al., 2014; S= Shirzadi et al., 2011 downregulated (x) in *cdka;1*-screen.

### Regulation of Parent-of-Origin Allelic Expression

We wanted to investigate the mechanisms involved in mediating imprinting on a larger scale. To this end we used mutants in epigenetic pathways previously demonstrated or hypothesized to be instrumental in setting up imprinting in the gametophyte germline (Figure 4). By using a crossing scheme that incorporates mutant lines as maternal or paternal contributors, we dissected the requirement of these epigenetic modifications for setting up imprinting in the affected parental allele. Crosses were made as described in Figure 4 and cleared whole-mount microscopy revealed that the tissue harvested had reached the same developmental stage in 4DAP seeds (SFigure 7; STable 7). cDNA libraries were prepared by the same procedure as for ecotype crosses described earlier (see Materials and Methods for details) and maternally and paternally derived reads were detected using the IRP as described previously (SData 9).

**Figure 4:**
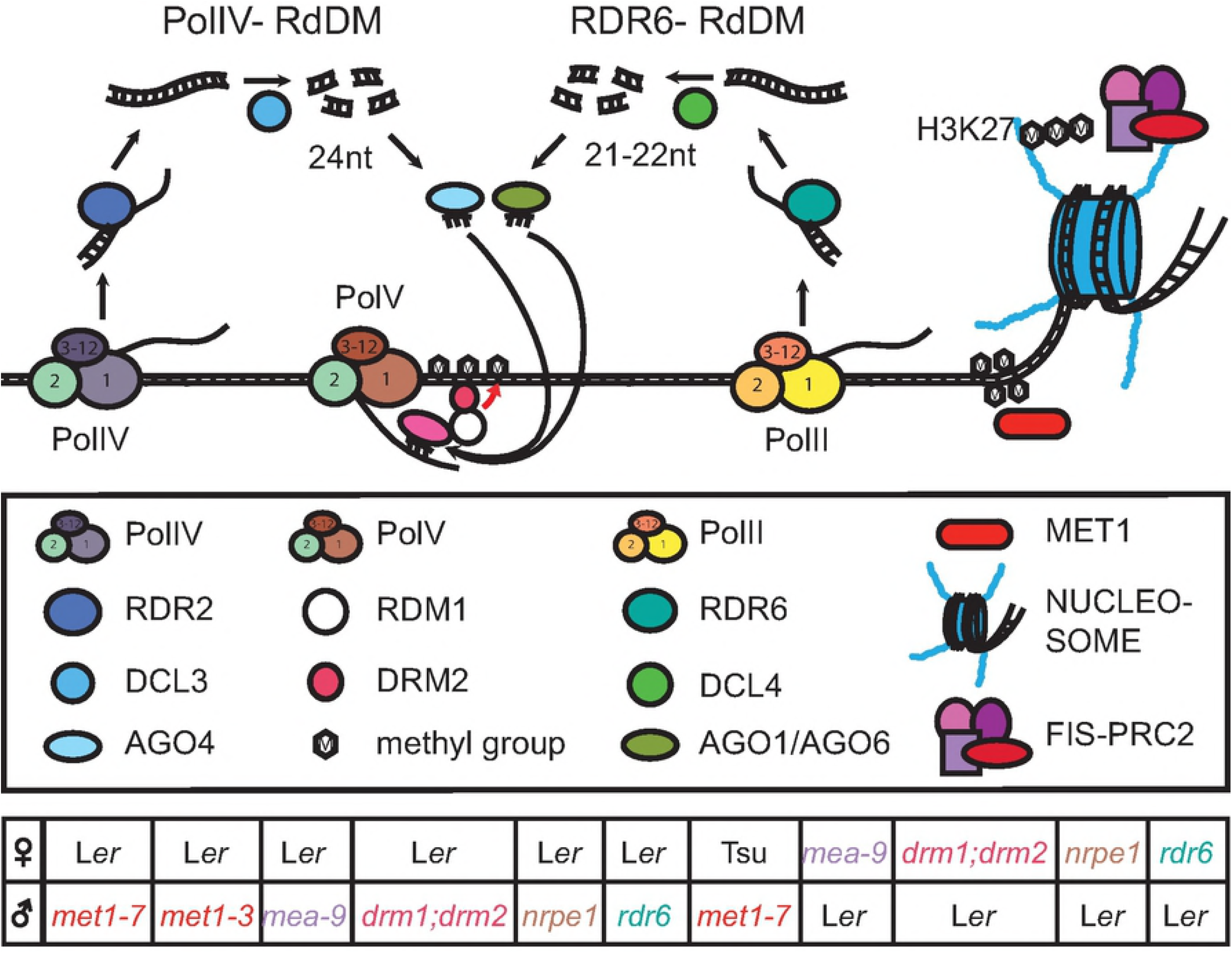
Summary of the epigenetic mechanisms postulated to govern the regulation of imprinted gene expression. Canonical (PolIV-RdDM) and non-canonical (RDR6-RdDM) utilize small RNA of different origin to mediate *de novo* DNA methylation in all sequence contexts as well as maintenance of CHH methylation. MET1 is the main methyltransferase responsible for maintenance of CG methylation. The FIS-PRC2 complex mediate trimethylation of the 27^th^ amino acid (Lysine) on the tale of histone 3 (H3K27-3me). Mutants (Col-0 background) of the various pathways investigated in this study are listed together with the wild type cross partner (L*er* or Tsu).

The mechanistic requirement of epigenetic regulators such as MET1 and MEA in establishment of genomic imprinting is traditionally demonstrated by change in maternal to paternal ratios in crosses where the epigenetic regulators are mutated in one parent. Generally, MET1 is hypothesized to silence the paternal allele of MEGs whereas MEA is repressing the maternal allele of PEGs (Moreno-Romero et al., 2016; Hsieh et al., 2011; Gehring and Satyaki, 2017). In order to analyze the overall effect of mutants of *MET1* and *MEA* in our dataset, we plotted parental ratios of identified MEGs and PEGs in crosses with paternal *met1* or maternal *mea-9* (SFigure 8, SData 9). Indeed, we could observe a ratio change towards biparental expression for MEGs in paternal *met1* crosses, whereas the PEGs in the same cross were mainly unchanged (SFigure 8A). In the maternal *mea-9* cross, the PEG ratios were changed towards biparental expression leaving MEGs more or less unchanged (SFigure 8B).

However, here we argue that a quantitative change in maternal to paternal expression ratio may not reflect a qualitative and specific expression change in the mutated parent. Changes in the wild type parent could also change the ratio and mimic a change in imprinting status. We have therefore analyzed changes in the mutated parent, i.e. maternal to maternal and paternal to paternal ratios between wild type and mutant crosses. To this end, we normalized all reads (RPM) to enable comparison of replicates from different crosses (SData 10). To evaluate the change of expression ratio in mutant parents we used the ratio of informative reads from the mutant parent versus the same parent in a wild type cross (e.g. L*er* x Col^mutant^ vs. L*er* x Col^wild type^). Furthermore, we assumed that deregulation of imprinted expression only takes place in the mutant parent. Consequently, the distribution of the wild type informative read ratios was used to infer the expected variation in the dataset. This distribution was used to create a two-sided prediction interval (Meeker et al., 2017) for the change in reads that is expected to be observed at a significance level of 0,05 (SData 10). The prediction interval was subsequently used to set a threshold to detect significant change in ratios of mutant parent reads to wild type parent reads. Observed mutant parent/wild type parent ratios outside of the prediction interval were identified as significantly changed. In a second step, to further account for variation between replicates that make up the average ratios, we performed a student’s t-test on the normalized read counts for the genes that were observed to have mutant parental ratios outside the prediction interval.

We compared expression from the paternal allele of MEGs in crosses where heterozygous *met1-7*^+/-^ and homozygous *met1-3*^-/-^ were used as pollen donors in crosses to WT L*er* or Tsu-1 mothers (Figure 5, SData 10). Many MEGs were notably changed in their paternal expression ratio whereas the maternal ratio was not significantly changed (Figure 5A, STable 8, p<0.05). As a control, PEGs were assessed from the same cross, and in contrast to MEGs, most PEG expression changes were within the two-sided prediction interval and hence did not change the true paternal ratio significantly (Figure 5B). More than 60 MEGs (25%) were significantly changed solely in paternal expression ratio in one or more *met1* cross combinations (STable 8, p<0.05). These comprised well-studied MEGs including *FIS2, FWA* and *MOP9.5*, but also novel MEGs from this study such as *GH9B8* (AT2G32990), *NAKR2* (AT2G37390) and *IQD16* (AT4G10640) (Table 2, STable 8). Except for the level of deregulation, there was not an obvious difference between heterozygous and homozygous *met1* fathers. Interestingly, some MEGs (5%) demonstrate significant paternal downregulation or downregulation from both genomes in these crosses, as exemplified by *MPC* and *AGL36* (Figure 5A, SData 10).

**Figure 5:**
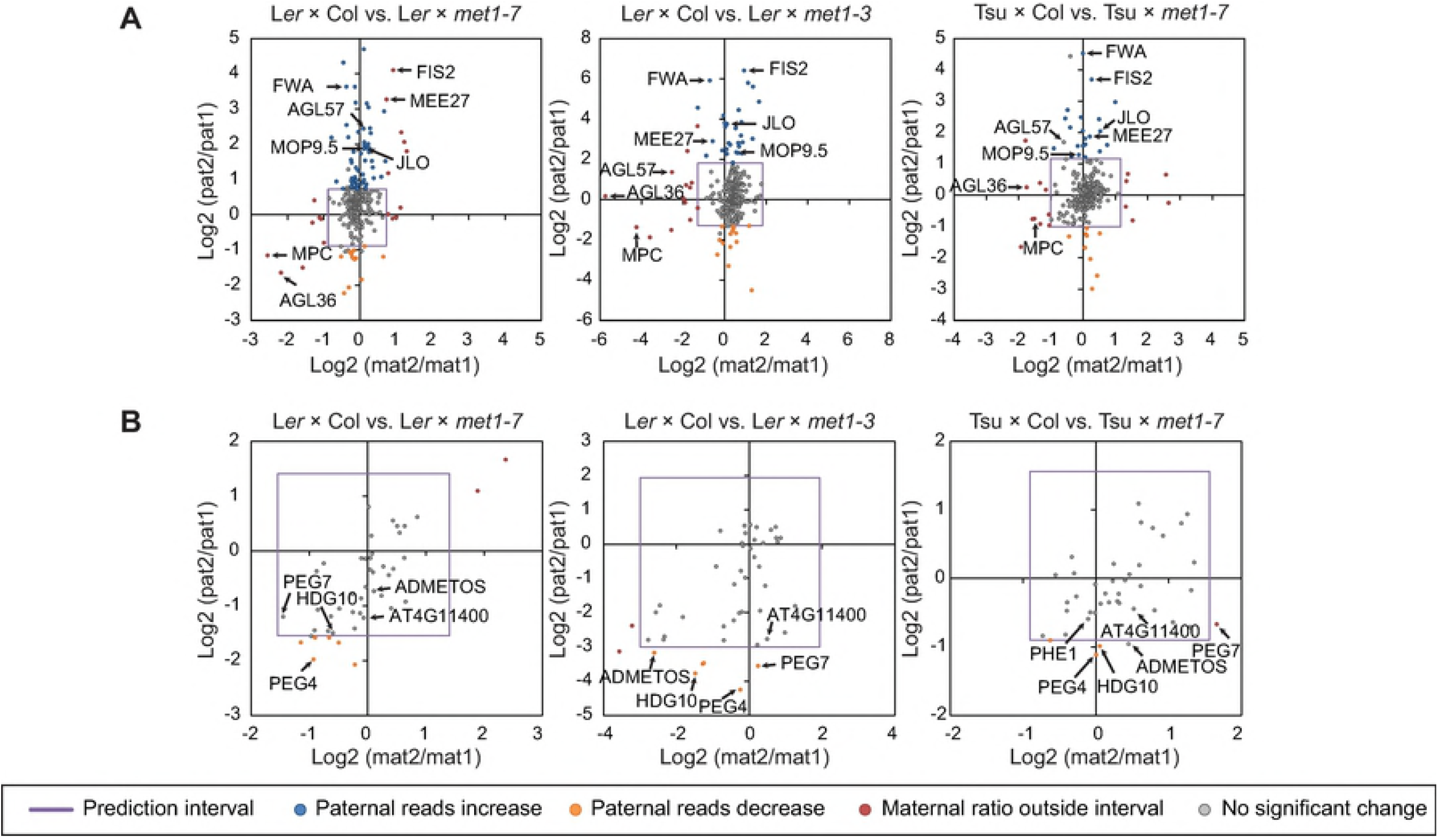
Impaired MET1 activity in pollen prior to fertilization mainly show upregulation of the paternal allele for maternally expressed genes (MEGs). Ratios of the change of maternal reads in mutant (mat2): maternal reads in wild type (mat1) was used to create a prediction interval (purple square) for analysis of paternal ratios; paternal reads in mutant (pat2): paternal reads in wild type (pat1). Based on the prediction interval made on maternal reads, genes where maternal reads fell outside the interval were not considered as significant (red circles). **A)** Several well-known MEGs show upregulation of the paternal allele in *met1*-mutants (blue circles). **B)** Paternally expressed genes are regulated in pollen to a lesser extent by MET1, although some genes show downregulation of the paternal allele (orange circles).

Next, we asked to what extent equal biparental expression levels were gained in crosses with heterozygous *met1* fathers. In a simplistic scenario (disregarding potential epimutations in *met1* mutant), half of the male gametes lack *MET1* activity and we assume that expression from the mutant paternal allele would become derepressed up to the level of the maternal alleles. Indeed, paternal expression levels up to around 50% can be observed, such as AT3G10900 and AT3G23060, but also low to intermediate level of upregulation of a few percent (Table 3). The latter cases still behave as MEGs even though the paternal expression is significantly elevated, suggesting that still other epigenetic mechanisms are repressing the paternal allele. We conclude that reactivation of biparental expression in MEGs by depletion of *MET1* is a quantitative modification rather than an on/off switch. Furthermore, the paternal expression levels were not observed to exceed maternal levels in a notable manner, suggesting a scenario where both parental alleles have the same potential level of expression. To our knowledge, no previous reports show imprinted allelic regulation in such a level of detail as described here.

**Table 3:** Selection of MEGs that show deregulation of paternal reads only, in the absence of MET1 paternal crossing partner

| AGI | Short name | Paternal reads |  | pat2/pat1<br>ratio | LxC<br>1/FC*100 | Lxm<br>1/FC*100 | Paternal reads |  | pat2/pat1<br>ratio | TxC<br>1/FC*100 | Txm<br>1/FC*100 | #<br>ecotypes |
| --- | --- | --- | --- | --- | --- | --- | --- | --- | --- | --- | --- | --- |
|  |  | LxC | Lxm |  |  |  | TxC | Txm |  |  |  |  |
| AT1G59930 | AT1G59930 | 6,0 | 52,7 | 7,64 | 3 % | 23 % | 14,4 | 63,0 | 4,15 | 7 % | 26 % | CLT |
| AT2G18880 | VEL2 | 1,9 | 15,9 | 5,83 | 3 % | 35 % | 6,1 | 23,9 | 3,49 | 9 % | 30 % | CLT |
| AT2G30590 | WRKY21 | 64,4 | 299,6 | 4,59 | 6 % | 37 % |  |  |  |  |  | CL |
| AT2G32990 | GH9B8 | 0,7 | 41,7 | 25,89 | 0 % | 18 % |  | Fail tests |  |  |  | CLT |
| AT2G34880 | MEE27 |  | Fail tests |  |  |  | 13,2 | 51,1 | 3,67 | 10 % | 33 % | CLT |
| AT2G34890 | AT2G34890 |  | Fail tests |  |  |  | 5,0 | 17,1 | 3,03 | 13 % | 28 % | CLT |
| AT2G35670 | FIS2 |  | Fail tests |  |  |  | 6,5 | 96,2 | 12,92 | 2 % | 24 % | CLT |
| AT2G37390 | NAKR2 | 0,3 | 14,9 | 12,38 | 0 % | 10 % |  | Fail tests |  |  |  | CLT |
| AT3G10900 | AT3G10900 | 17,2 | 61,5 | 3,43 | 15 % | 48 % |  | Fail tests |  |  |  | CLT |
| AT3G23060 | AT3G23060 | 9,3 | 46,1 | 4,55 | 7 % | 59 % | 15,7 | 90,3 | 5,46 | 7 % | 60 % | CLT |
| AT3G51895 | SULTR3;1 | 8,5 | 85,6 | 9,07 | 0 % | 5 % | 23,1 | 106,7 | 4,46 | 1 % | 3 % | CLT |
| AT4G00220 | JLO | 2,6 | 12,3 | 3,71 | 1 % | 2 % | 2,2 | 12,3 | 4,10 | 0 % | 1 % | CLT |
| AT4G10640 | IQD16 | 36,8 | 129,2 | 3,45 | 4 % | 16 % |  | Fail tests |  |  |  | CLT |
| AT4G21430 | B160 | 7,3 | 26,7 | 3,34 | 2 % | 8 % |  | Fail tests |  |  |  | CLT |
| AT4G25530 | FWA | 10,6 | 142,3 | 12,41 | 1 % | 21 % | 5,4 | 146,9 | 23,07 | 1 % | 22 % | CLT |
| AT5G07460 | PMSR2 | 15,3 | 80,0 | 4,97 | 7 % | 35 % |  | Fail tests |  |  |  | CLT |
| AT5G24240 | MOP9.5 | 81,7 | 301,2 | 3,65 | 4 % | 14 % | 131,8 | 321,7 | 2,43 | 5 % | 13 % | CLT |
| AT5G37560 | AT5G37560 | 0,4 | 27,8 | 19,94 | 1 % | 58 % |  | Fail tests |  |  |  | CLT |
| AT5G41830 | AT5G41830 | 2,0 | 23,7 | 8,25 | 1 % | 13 % | 1,7 | 13,3 | 5,37 | 7 % | 39 % | CLT |
| AT5G48650 | AT5G48650 | 83,2 | 460,6 | 5,48 | 4 % | 16 % |  | Fail tests |  |  |  | CLT |

Average normalized paternal read counts for *Ler*Col (LxC) and *Tsux*Col (TxC) crosses compared to the respective *met1-7* mutant cross (Lxm and Txm), and the associated ratios. For all read counts reported, a student's t-test showed significant change in mutant compared to WT (p-values <0,05). As an expression of how much the paternal allele is expressed compared to the maternal, fold change (FC) values from differential expression of maternal and paternal alleles was used (1/FC). Some genes fail the prediction interval test (indicated by fail tests) because the maternal reads are outside the initial prediction interval, paternal reads from these were not analysed. CLT = SNPs in all ecotypes tested, CL= only SNP between Col-0 and *Ler*.

The PRC2 member MEA is a key player for the establishment of silencing of the maternal allele of PEGs (Köhler et al., 2004; 2003; Makarevich et al., 2008). We compared expression from the maternal allele of PEGs in crosses where heterozygous *mea-9*^+/-^ mothers were crossed to WT L*er*. (SData 10). Indeed, depletion of maternal *MEA* strongly deregulates most PEGs (78%) (Figure 6A). When maternal *MEA* is mutated, all significant expression change occurs from the maternal allele (Figure 6A), and with the majority of PEGs (69% of deregulated PEGs), maternal allele reactivation and elevated expression levels are observed (Table 4). This includes most previously documented PEGs such as *HDG10, MEO* and *AGL23*. In contrast, the majority of MEGs are within the prediction interval, and thus not significantly deregulated in the same cross (Figure 6A, B). As expected, the same situation is found for both MEGs and PEGs in the reciprocal mutant cross (Figure 6C, D).

**Table 4:** PEGs regulated by MEA as shown by deregulation of the maternal allele in *mea-9*xL*er*.

| AGI | Gene name | Short description | maternal reads |  | MAT2/<br>MAT1 | ColxLer<br>1/FC*100 | *mea-9xLer*<br>1/FC*100 |
| --- | --- | --- | --- | --- | --- | --- | --- |
|  |  |  | ColxLer | *mea-9xLer* |  |  |  |
| AT1G23320 | TAR1 | tryptophan aminotransferase | 0,0 | 14,0 | 14,63 | 1 % | 48 % |
| AT1G34650 | HDG10 | homeodomain GLABROUS 10 | 9,8 | 119,0 | 11,09 | 5 % | 46 % |
| AT1G48910 | YUC10 | Flavin-containing monooxygenase | 10,8 | 121,5 | 10,40 | 10 % | 80 % |
| AT1G49290 | PEG2 | Rho guanine nucleotide exchange | 13,8 | 95,7 | 6,53 | 4 % | 19 % |
| AT1G60400 | PEG3 | F-box/RNI-like superfamily protein | 5,2 | 44,2 | 7,26 | 5 % | 46 % |
| AT1G65360 | AGL23 | AGAMOUS-like 23 | 8,7 | 80,7 | 8,41 | 6 % | 79 % |
| AT1G66630 | PEG4 | Protein with RING/U-box | 25,9 | 105,1 | 3,94 | 15 % | 53 % |
| AT1G67820 | AT1G67820 | Protein phosphatase 2C | 117,1 | 649,3 | 5,51 | 7 % | 88 % |
| AT1G67830 | FXG1 | alpha-fucosidase 1 | 383,5 | 2217,2 | 5,77 | 21 % | 76 % |
| AT2G20160 | MEO | E3 ubiquitin ligase SCF complex | 41,4 | 673,6 | 15,92 | 19 % | 74 % |
| AT2G25700 | SK3 | SKP1-like 3 | 91,0 | 504,2 | 5,49 | 34 % | 87 % |
| AT2G32370 | HDG3 | homeodomain GLABROUS 3 | 237,4 | 1722,4 | 7,23 | 24 % | 39 % |
| AT2G36560 | PEG5 | AT hook motif DNA-binding | 50,1 | 257,9 | 5,07 | 19 % | 98 % |
| AT3G49770 | PEG6 | hypothetical protein | 0,4 | 18,8 | 14,27 | 1 % | 38 % |
| AT3G50720 | PEG7 | Protein kinase superfamily protein | 1,3 | 28,2 | 12,92 | 0 % | 6 % |
| AT4G10160 | AT4G10160 | RING/U-box superfamily protein | 7,9 | 73,1 | 8,29 | 7 % | 48 % |
| AT4G11400 | AT4G11400 | ARID/BRIGHT DNA-binding | 42,6 | 247,9 | 5,71 | 28 % | 85 % |
| AT4G23110 | AT4G23110 | insulin-like growth factor | 14,2 | 65,0 | 4,34 | 10 % | 98 % |
| AT5G15140 | PEG9 | Galactose mutarotase-like | 5,1 | 37,7 | 6,36 | 15 % | 50 % |
| AT5G50470 | NF-YC7 | nuclear factor Y, subunit C7 | 8,8 | 104,4 | 10,73 | 4 % | 84 % |

Average normalized maternal reads for ColxLer compared to mea9xLer and associated ratio, show a high upregulation of the maternal allele for most PEGs in mutant cross. As an expression of how much the paternal allele is expressed compared to the maternal, fold change (FC) values from differential expression of maternal and paternal alleles was used (1/FC).

**Figure 6:**
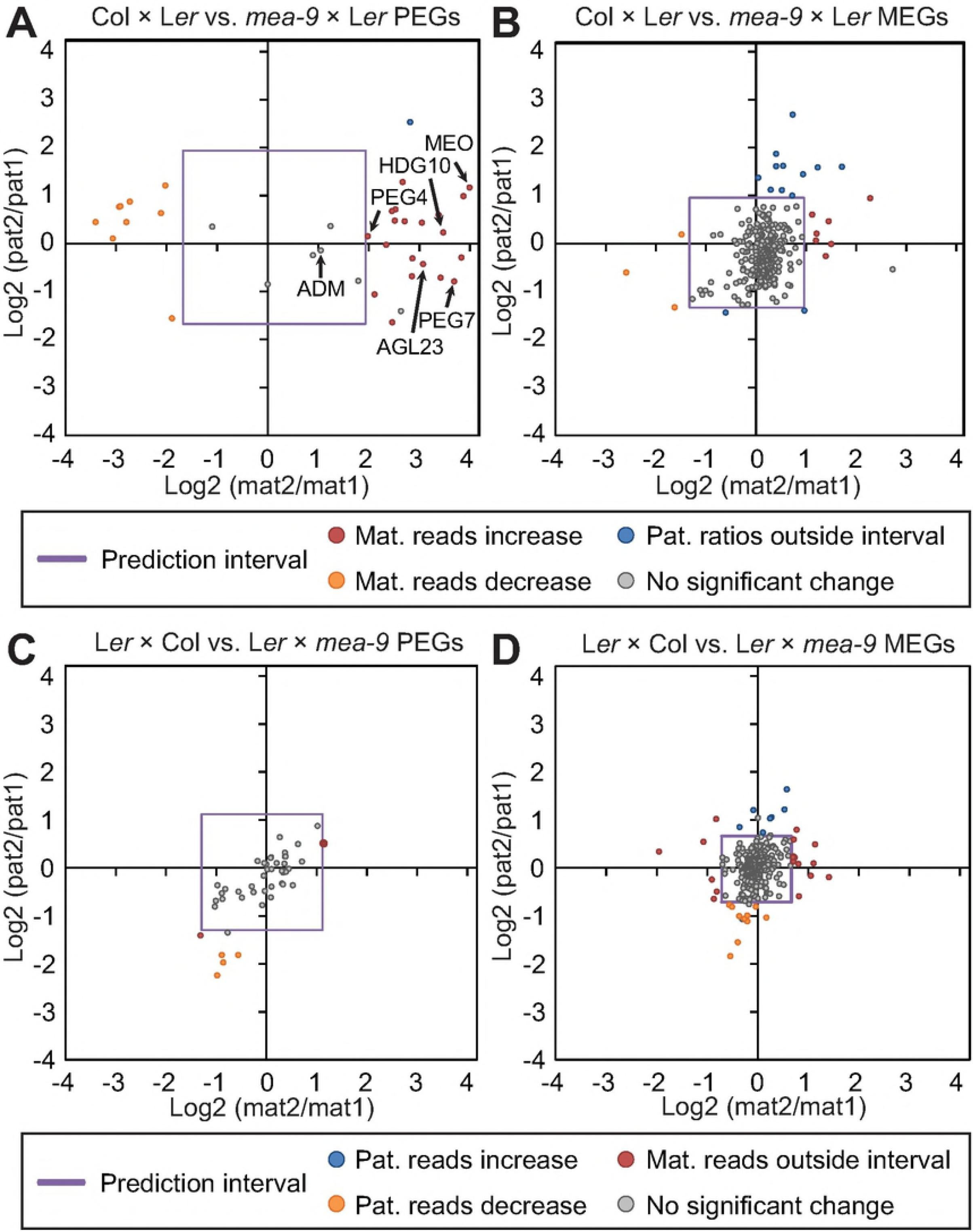
Maternal reads are upregulated for paternally expressed genes (PEGs) when PRC2 activity is impaired on the maternal side. Ratios of the change of paternal reads in mutant (pat2): paternal reads in wild type (pat1) was used to create a prediction interval (purple square) for analysis of maternal ratios; maternal reads in mutant (mat2): maternal reads in wild type (mat1). **A)** Several well-known PEGs show upregulation of the maternal allele in *mea-9*xL*er*. **B)** Maternally expressed genes are not greatly affected by impaired PRC2 activity in *mea-9*xL*er*. **C)** Regulation of PEGs in L*er*x*mea-9*. **D)** Maternally expressed genes are not greatly affected by impaired PRC2 activity in L*er*x*mea-9*. Based on the prediction interval made on paternal reads, genes where paternal reads fell outside the interval were not considered as significant; blue circles in **A)** and **B)**, red circles in **C)** and **D)**.

We concluded that the extent of equal parental expression in crosses with heterozygous *mea-9* mothers was close to bi-allelic. With few exceptions, the expression of the maternal alleles was at the same as, or higher level than the paternal expression (Table 4). We also observed that the maternal alleles of some deregulated PEGs (31%) were significantly downregulated in the same cross, suggesting that imprinting of these genes is only indirectly affected by MEA (Figure 6A). Furthermore, a handful of MEGs had moderately elevated paternal expression or appeared to turn into stronger MEGs (Figure 6B). This may suggest that MEA can play an indirect role or act in post-fertilization regulation of MEGs as demonstrated previously (Shirzadi et al., 2011).

### Most Imprinted Genes Are Not Regulated by MET1 or PRC2 MEA

In the previous experiments, we have verified establishment of imprinting patterns by *MET1* and *PRC2 MEA*. The majority of PEGs show restored maternal expression in maternal *PRC2 mea-9* mutant crosses, indicating that PRC2 is a major determinant of repressing the maternal allele of PEGs. However, for the majority of MEGs, depletion of *MET1* did not lead to restored expression of the paternal allele (Figure 7, STable 9). For identified MEGs that were >10-fold maternally biased (SData 7), only one third demonstrated paternal reactivation upon mutational removal of paternal *MET1* (Figure 7A, STable 9A). One third of the verified PEGs showed significant downregulation of the paternal allele in the same cross (Figure 7B, Figure 5B), hypothetically involving a DNA methylation dependent regulation mechanism of MEA activity (Makarevich et al., 2008; Köhler et al., 2012; Weinhofer et al., 2010). Two thirds of the PEGs could be verified to have significantly elevated expression from the maternal allele in the maternal absence of *MEA* (Figure 7C, STable 9B), as opposed to only a minor effect on MEGs (Figure 7D). Taken together, less than half of the unambiguously imprinted genes in this study can be assigned a major mechanism for imprinting. We therefore investigated if the RdDM pathway (Figure 4) could be instrumental in setting up imprinting patterns as suggested previously (Vu et al., 2013; Satyaki and Gehring, 2017).

**Figure 7:**
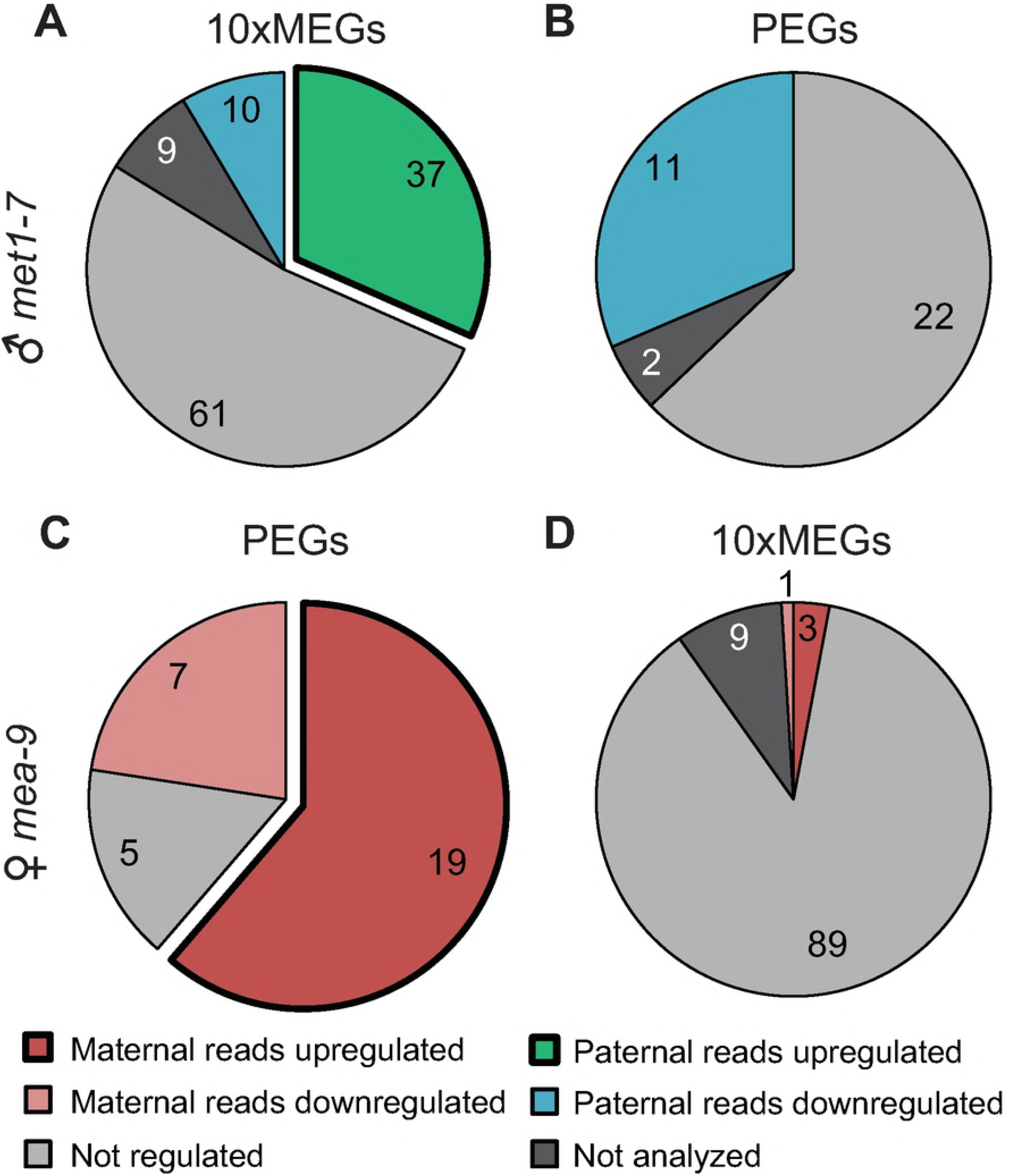
Only a fraction of maternally- and paternally expressed genes (MEGs and PEGs) are regulated by MET1 or MEA respectively. **A)** and **B)** More than half of identified 10x MEGs and PEGs show no deregulation in the absence of MET1 in pollen (♂ *met1-7*). **C)** Most PEGs are regulated by MEA and **D)** very few MEGs are deregulated when *mea-9* is used as the maternal cross partner.

### RdDM Alone Is Not a Major Mechanism for the Establishment of Genomic Imprinting

We compared imprinted expression from maternal and paternal alleles of MEGs in reciprocal crosses between WT L*er* and homozygous mutants of *DRM1;DRM2, NRPE1(PolV)* and *RDR6* (Figure 4, SData 10). Compared to *met1* and *mea-9* crosses the effect of mutation of RdDM produce less variation and deregulation of imprinted genes (Figure 8, cf. Figure 5). Hypothetically, in the cross where RdDM is blocked in the male cross partner, we would expect the paternal genome to be upregulated if RdDM is involved in silencing of the paternal allele in MEGs. In general we find less upregulated genes and a modest fold change ratio (Figure 8). A few genes are however significantly regulated outside the prediction interval but many of these do not pass a two-sided t-test for significant difference between paternal allele expression. In order to lend higher reliability to our findings we have used crosses with several mutants affected in both canonical and non-canonical RdDM. In canonical RdDM, *drm1;drm2* and *nrpe1* mutants are expected to display deregelation of the same targets. In non-canonical RdDM both of the latter and also *rdr6* should be implicated. Comparing upregulated paternal alleles of MEGs in the WT x RdDM-mutant cross a total of 21 genes showed this effect in one or more of the mutants used (Figure 8). Two targets overlapped between crosses with *drm1;drm2* and *nrpe1* but not *rdr6*, and may be specific to the canonical pathway (Figure 8D). No targets overlapped in all mutant combinations, representing the non-canonical pathway. Two overlapping targets between *rdr6* and *nrpe1* were found, but could not be supported by *drm1;drm2*, thus may suggesting that another DNA methyltransferase is involved. This is in line with and confirms previous reports suggesting that RdDM is not active in in sperm cells and absent after fertilization prior to 4DAP and endsperm cellularization (Borges et al., 2008; Calarco et al., 2012; Moreno-Romaro et al., 2016; Ingouff et al., 2017). Since homozygous mutants were used in this experiment, a lack of contribution of RdDM through male sporophytic tissue is not sufficient to deregulate imprinting.

**Figure 8:**
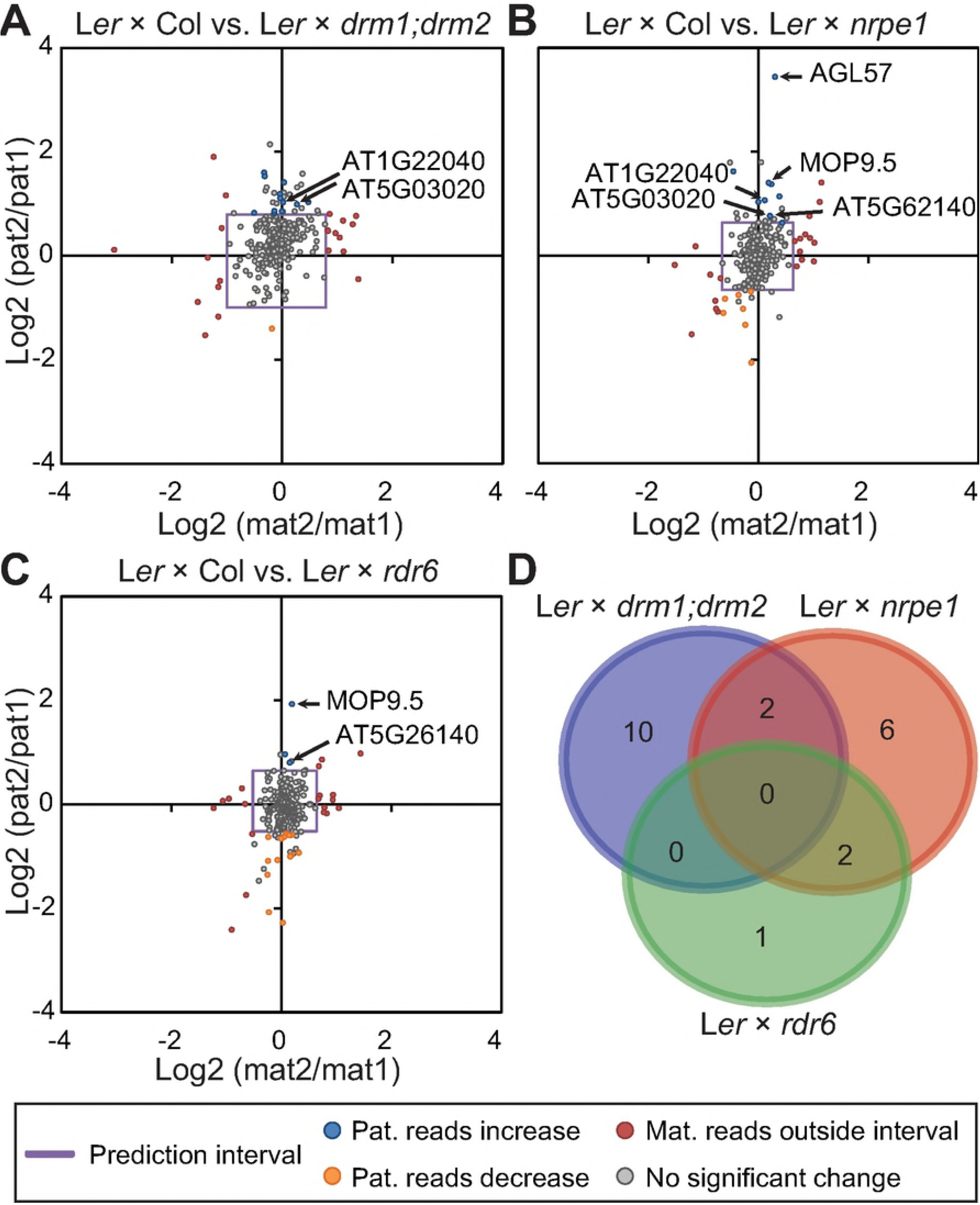
Mutants in RdDM are not sufficient to deregulate imprinted gene expression of maternally expressed genes (MEGs). Ratios of the change of maternal reads in mutant (mat2):maternal reads in wild type (mat1) was used to create a prediction interval (purple square) for analysis of paternal ratios; paternal reads in mutant (pat2): paternal reads in wild type (pat1). Based on the prediction interval made on maternal reads, genes where maternal reads fell outside the interval were not considered as significant (red circles). Low degree of deregulation is observed of the paternal allele for MEGs in crosses with mutants of the RdDM pathway **A)** L*er*x*drm1;drm2* **B)** L*er*x*nrpe1*, **C)** L*er*x*rdr6*. **D)** Limited overlap of the few deregulated genes in the RdDM mutant crosses of **A)**, **B)** and **C)**. Overlapping genes in **D)** are indicated in the Log2-plots.

Although RdDM is not present in early endosperm (Moreno-Romero et al., 2016) we tested the hypothesis that CHH marks set by RdDM is involved in expression of the paternal genome yielding reduced paternal expression after fertilization, analogous to previously shown for CG methylation (Makarevich et al., 2008). In our dataset we find 11 downregulated targets of which 3 are shared by *drm1;drm2* and *nrpe1* (SFigure 9D). The *rdr6* cross did not generate any deregulation of the paternal allele of PEGs.

To inspect the effect of RdDM in maternal gametes we also performed crosses with the mutant as maternal cross partner. For MEGs, we tested the hypothesis that RdDM is directly or indirectly involved in maintenance or up-holding expression from the maternal allele. In such a scenario, loss of RdDM would result reduced maternal expression if DNA methylation is a means to avoid silencing. Only a weak downregulation effect can be observed whereas the effect on upregulation was more prominent, indicating a silencing role of RdDM (SFigure 10).

In the same manner we analyzed PEGs when RdDM is depleted in the mother. Here, we hypothesized that RdDM is required to keep the maternal allele silent and that in the mutants we get upregulation of the maternal allele. All crosses taken together only produced one gene slightly (2-fold) upregulated. We detected, however, a downregulation effect on several genes in the canonical RdDM mutants (SFigure 11).

## Discussion

### Conservation of Imprinting in *A. thaliana* Ecotypes

We have used sequence capture technology in combined with high throughput sequencing to investigate imprinted allelic expression at great sequence depth. In the same manner, epigenetic mechanisms that are responsible for setting and maintaining the imprinting pattern were systematically investigated. In order to limit the number of genes and reach sufficient read-depth, we captured transcripts from previously reported imprinted gene panels (Wolff et al., 2011; Gehring et al., 2011; Hsieh et al., 2011) for which there was little overlap between panels and a dataset based on the functional absence of a paternal genome in the endosperm (Shirzadi et al., 2011; Nowack et al., 2006; Aw et al., 2010). Using three different accessions of *A. thaliana*; Col-0, L*er* and Tsu in reciprocal crosses enabled the detection of parental transcript origin in the developing seed at high sequencing depth to give robust data for analysis of variation between ecotypes, but also to ensure high sensitivity for genes with very low expression.

We developed a bioinformatics pipeline referred to as the Informative Reads pipeline (IRP) to detect parent-of-origin allele specific expression based on all allele specific sequence polymorphisms. Imprinted genes could confidently be identified from reciprocal crosses between Col-0 and L*er* and between Col-0 and Tsu. An important consideration was maternal seed coat expression in the sequencing data obtained. Due to RNA extraction from whole seed, maternally derived tissue such as seed coat could contribute to the detection of false positive MEGs. We therefore filtered away potential seed coat enriched genes based on expression profiles for the different seed compartments (Belmonte et al., 2013) but retained genes with seed coat expression not exceeding endosperm expression levels. The remaining 282 candidate MEGs and 35 candidate PEGs show parent-of-origin specific expression in all three ecotypes.

From the list of candidate MEGs, 30 genes not previously verified as imprinted (Shirzadi et al., 2011) were identified as having more than 10-fold maternal expression compared to paternal expression. Even though previous filtering methods based on available microarray data are controversial (Schon and Nodine, 2017), we argue that our developed threshold to define seed coat expression is sufficient to determine imprinting. Previously, Schon and Nodine required an 8-fold expression of a transcript in a given seed compartment compared to other seed compartments to be significantly determined as specifically enriched. In our setup, we exclude all targets with more than 1-fold expression in seed coat, and therefore our dataset is devoid of targets that have higher expression in the seed coat than other compartments. This is argued by the fact that if a gene is equally expressed in endosperm and seed coat it may still be imprinted in the endosperm. Thus, the high expression levels observed for some of the novel MEGs reported here is not likely a product of the maternal seed coat as it is not substantially detected in the seed coat from analysis using the above discussed microarray data. Here, we reason that in a biological context, genomic imprinting in the endosperm *per se* does not depend on the absence or presence of expression in other seed tissues. Albeit seed coat expression indeed represents a technical limitation, we argue that the filtering method applied for identification of parentally biased expression presented by our moderate stringency dataset allows unambiguous identification of imprinted genes.

Identification of false positive PEGs do not suffer from contamination of maternally derived tissues in the same way as MEGs, however PEGs may not be detected due to maternal seed coat expression masking biased paternal expression. 35 PEGs were reciprocally detected in at least two ecotypes and of these, only one gene had not previously been reported as a PEG. Taken together this confirms the predominance of MEGs in comparison to PEGs in *A. thaliana*. Even though our 1011 targets are selected based on previous investigations, which indeed are overrepresented by MEGs (Wolff et al., 2011; Shirzadi et al., 2011; Gehring et al., 2011; Hsieh et al., 2011), the genes included from the Shirzadi et al. (2011) study was initially designed as a screen to identify paternally expressed genes and thus should offer potential to discover novel PEGs. The overrepresentation of imprinted MEGs compared to imprinted PEGs supports a co-adaptation theory that explains how imprinting and maternally biased expression evolve. This theory has been supported by the fact that the offspring does not necessarily benefit from unlimited supply of energy resources from the mother, as big size does not necessarily increase fitness of the offspring (Wolf and Hager, 2006).

### Genomic Imprinting Is a Quantitative Phenomenon

In our analysis of parentally biased expression we find that most genes are not strictly mono-allelic expressed and even though activity from only one allele is more prominent for PEGs, our data set suggests that maternal and paternal allelic expression is differentially regulated and that the regulation is not a binary on/off situation. This result is also supported by previous investigations of imprinting (Gehring et al., 2011, Hsieh et al., 2011), and is in support of the differential dosage hypothesis, which states that an allele can be partly silenced if the obtained expression level is more optimal for that gene product, and that complete silencing of one allele is not necessarily the most optimal (Dilkes and Comai, 2004).

Gehring and Satyaki (2017) suggest that partial imprinting can arise in two scenarios. Either by partial imprinting within each cell, or as a mixture of expression patterns between cells in the endosperm. Previous reports as well as our study are not sensitive enough to detect different expression patterns in sub-compartments of the endosperm. A gene might be imprinted with a strong parental bias in one part or time point within the endosperm and less or even without parental bias in another part or time point within the endosperm. Furthermore, partial imprinting suggests that the co-adaptation theory may provide a stronger selective pressure on the maternal side compared to the paternal side, while the differential dosage hypothesis can explain parental biased expression in a wide variety of genes and not only genes targeting nutrient allocation.

### MET1 and PRC2 Insufficiently Explain the Regulation of Imprinting

DNA methylation maintained through MET1 and histone modification by the PRC2 are the two main epigenetic modifications previously shown to affect imprinted genes. In general, for maternally expressed genes, it is the DNA glycosylase DME that mediate expression of the maternal allele, however, MET1 is required to keep the paternal allele silent. For PEGs, it is histone methylation by the PRC2 which establishes silencing of the maternal allele. Also in this case, it has been proposed that DNA methylation is required in some part to mark the paternal allele, but in this setting to avoid silencing by PRC2 (Makarevich et al., 2008).

Although efforts have been made to systematically investigate the effect of these mechanisms on imprinted genes (Hsieh et al., 2011; Wolff et al., 2011), previous individual studies suffer from limited overlap between the sets of imprinted genes. Even though ratios between maternal and paternal reads in wild type have been compared to ratios in mutants, ratios between maternal and paternal reads may not always reflect the expected change. In other words, a decrease in maternal:paternal ratio, may be due to maternal expression decreasing or paternal expression increasing. Similarly, the ratio may seemingly not have changed if maternal and paternal reads have experienced the same decrease or increase.

Two different *met1* alleles (homozygous *met1-3*, heterozygous *met1-7*) were used to investigate the effect of MET1 on the paternal genome in establishing imprinting. By means of differential expression analysis of maternal to paternal reads, differences in fold change i.e. maternal:paternal ratio could be observed in the L*er* x *met1-3* compared to L*er* x Col WT cross. As expected by the number of gametes affected, these differences were stronger in the crosses using the homozygous compared to crosses using the heterozygous *met1*- alleles. This is however, not a strong argument since epimutations may occur in the homozygous mutant. Nonetheless, it is a major point that we indeed observe a gametophytic requirement and effect of depletion of MET1.

On the other side, mainly PEGs were affected by impaired PRC2 activity on the maternal side comparing *mea-9* x L*er* and Col x L*er* crosses. To further explore the cause of the FC differences and to also get a more correct picture of how these mutants affect imprinted genes, the actual change of maternal only reads or paternal only reads was compared between mutant and WT crosses. This was analyzed by creating a prediction interval (Meeker et al., 2017). In general, in a mutant cross, the reads originating from the WT parent are not expected to be directly affected by the mutation in the crossing partner. Therefore, the change in reads originating from the WT parent in a mutant cross were compared to reads from the same parent in the WT cross. E.g. in L*er* x *met1*, the L*er* reads were compared to L*er* reads in L*er* x Col creating a maternal:maternal ratio in this case. The distribution of these ratios were analyzed to define a 95% prediction interval in which expected variation is most likely to be found. The prediction interval was then applied to analyze the reads originating from the mutant parent compared to the same parental reads in the WT cross. In the case of L*er* x *met1*, this was the paternal mutant to paternal WT ratio, and only deviations outside the interval were considered true changes in allelic bias caused by the mutant. Furthermore, to consider also the variation between biological replicates a t-test was applied independently on reads from each replicate as the prediction interval was based on average read counts across biological replicates.

By applying this methodology, more than 60 genes displaying maternally biased expression in WT were identified as being significantly deregulated in one or several of the L*er* x *met1* crosses. These genes displayed a significant upregulation of the paternal allele when MET1 activity is impaired in pollen prior to fertilization, among these *FWA* and *FIS2* that previously have been shown to depend, directly or indirectly, respectively on MET1 silencing of the paternal allele (Kinoshita et al., 2004; Jullien et al., 2006; Wöhrmann et al., 2012).

In support of the idea that imprinting does not rigidly lead to mono-allelic expression, the analysis of reactivation of the paternal allele in the *met1* crosses show that some paternal alleles, but not all, are reactivated to the same level as the non-silenced allele. This suggests that methylation by MET1 does not necessarily act as a dual switch, but rather have the possibility to create a fine tuned expression level, possibly also in concert with other epigenetic regulation mechanisms.

Except for the level of deregulation, both heterozygous and homozygous *met1* fathers affected globally the same genes. Among those, some MEGs demonstrate significant paternal downregulation or downregulation from both genomes, as exemplified by the well-studied imprinted genes *MPC* and *AGL36* (Shirzadi et al., 2011), suggesting that previously reported biallelic expression in *AGL36* in lack of paternal MET1 is rather an effect of maternal *AGL36* downregulation than upregulation of the paternal *AGL36* allele, i.e. no direct effect on *AGL36*.

We conclude that reactivation of biparental expression in MEGs by depletion of *MET1* is a quantitative modification rather than an on/off switch. Furthermore, the paternal expression levels were not observed to exceed maternal levels in a notable manner, suggesting that structural variants of the two parental alleles do not interfere with transcriptional regulation and hence both alleles have the same expression potential. Consequently, differences in expression between both alleles in the wild type situation are of epigenetic origin.

Investigations of the PRC2 component MEA, showed an almost exclusive deregulation of paternally biased genes in crosses using maternal *mea-9* mutants. Prediction interval analysis showed that nearly all PEGs analyzed were affected in the *mea-9* mutant cross, and that in the majority of these, maternal transcripts were upregulated in line with the postulated function of PRC2 in the central cell. In contrast to MET1, MEA activity is important for the silencing of the maternal allele for most, if not all, PEGs, while MET1 clearly is not responsible for the repression of the paternal allele of majority of MEGs.

### RdDM Alone Is Not Required for Establishment of Imprinting

Using the established setup, the parental origin of expression of imprinted genes was also analyzed in crosses involving mutants of the RdDM pathway. Three different mutant lines were used in reciprocal crosses to L*er* to inspect potential maternal as well as paternal deregulation of imprinted genes. The *drm1;drm2* double mutant and *nrpe1* are both central actors of RdDM and both the canonical and non-canonical pathways depend on the NRPE1 subunit of PolV, and the DRM2 DNA MTase for targeting and methylation of DNA. The *rdr6* mutant was also included with the goal to distinguish the canonical and non-canonical pathways as RDR6 is involved in processing of RNA from different sources than in the canonical PolIV-dependent pathway.

MEGs have previously been shown to be enriched for paternally derived siRNA (Pignatta et al., 2014). Furthermore, a 5% general increase of maternal mRNA was observed in crosses involving the RdDM PolIV mutant *nrpd1*, and it was suggested that the silenced allele of imprinted genes is associated with siRNA (Erdmann et al., 2017).

In our data, and in strong contrast to the deregulation observed in *met1* and *mea-9* crosses, a limited deregulation was detected for imprinted genes in the crosses with disrupted RdDM activity. In accordance with Erdmann et al., although not addressing the regulation of the silenced allele, we do observe derepression of maternal mRNA from MEGs in crosses with paternal as well as maternal depletion of RdDM components. The observed effect is in line with previous observations (Erdmann et al., 2017), however limited compared to the effect observed for *met1* and *mea-9*.

RdDM is not active in sperm cells (Calarco et al., 2012) and in a hypothetical scenario where RdDM is involved in silencing of MEGs paternal alleles, this effect must be executed in the pollen vegetative cell or in the sporophyte. In this scenario, in paternally contributed RdDM mutants, we expect that paternal alleles may show patterns of reactivation. In the individual crosses with a homozygous RdDM mutant as the paternal contributor, a total of 21 genes show significant deregulation in one or more crosses with only little overlap between the different mutant crossing partners.

Nevertheless, one of the candidate MEGs to require RdDM activity from the male cross partner to keep the paternal allele silenced is *MOP9.5*. The paternal allele of *MOP9.5* shows a modest, but significant degree of reactivation in crosses with both *rdr6* and *nrpe1* as father in the cross. This may suggest that *MOP9.5* is regulated by a non-canonical RdDM pathway, however, in the *drm1;drm2* mutant cross the paternal allele shows no sign of deregulation, suggesting the observed regulation to be an indirect effect. *MOP9.5* is one of very few genes previously shown to be regulated by the RdDM pathway (Vu et al., 2013). In addition, this research team could show reactivation of the paternal allele in the *drm1;drm2* double mutant, but also when using the *nrpd2* RdDM mutant, affecting both, PolIV and PolV complexes. However, these mutant alleles cannot be used to discriminate between canonical and non-canonical pathways because PolV, DRM1 and DRM2 are postulated to be involved in all RdDM pathways. Vu et al. also studied the imprinting and regulation of *SDC*, which was also shown to be regulated by RdDM. In the filtering steps used in our study, *SDC* was excluded from the moderate stringency group because it had no expression information in the LCM data (Belmonte et al., 2013). SDC is therefore detected as MEG in the low stringency group, and also shows strong upregulation of the paternal allele in crosses with the *met1* mutants. In contrast to Vu and co-workers, we did not detect any *SDC* deregulation in crosses with RdDM mutants.

As expected, only very few imprinted genes show to be regulated solely by RdDM compared to MET1 and MEA as described above, and still the bulk of MEGs have not been connected to any regulatory mechanisms. Redundancy may be an issue regarding RdDM, but two of the common components for canonical PolIV-RdDM and the non-canonical RDR6-RdDM pathways suggested to date have been thoroughly investigated in the data presented here. These data show that RdDM does not contribute to a large extent to regulate imprinted genes prior to fertilization on either parental sides. It is possible that the RdDM machinery is required after fertilization, but considering the data presented here it is not likely an important regulator in stages up to 4 DAP. Collectively these results point to the fact that still other mechanisms and pathways for maintaining imprinting, especially for MEGs remains to be discovered.

The fact that siRNA map to imprinted genes point towards a connection to imprinting (Calarco et al., 2012). RdDM has been widely studied in relation to silencing of TEs; several alternative pathways have been shown to also exist for the non-canonical pathway, also those that are not dependent on the RDR6 polymerase investigated here (Teixeira et al., 2009). Even though all RdDM pathways are postulated to be targeted to the PolV scaffold, where DRM2 is the main MTase mediating DNA methylation in RdDM, DNA MTases may act redundantly.

### Concluding Remarks

Here we provide compelling evidence that PRC2 histone modification and DNA methylation by MET1 are only in part explaining the regulation of imprinting, as demonstrated by our finding that *most* parent-of-origin expressed genes are not derepressed in crosses with mutant *mea-9* and *met1*. Furthermore, we show that siRNAs through the RdDM pathway is likely not a regulator of imprinting to the level of MET1 and PRC2.

The data presented here suggest a current lack of a regulative mechanism explaining the majority of imprinted genes. A possible hypothesis to account for this is an interplay between different epigenetic pathways, that require a more advanced experimental setup to be verified. An example of this may be the downregulation of the paternal allele of PEGs in the absence of MET1 in pollen (Figure 5B, STable 10). A similar regulation is found to affect some of the same PEGs in crosses with paternal RdDM mutant partner (Figure 8, STable 10). Most of these common targets are also targeted in crosses with maternal *mea-9* (Figure 6, STable 10), and may suggest a common mechanism involved. However, other loci are not MEA targets and thus call for another mechanism to explain imprinting.

In a wider perspective, our findings give support to several of the theories explaining the evolution of imprinting. It has been postulated, that imprinted genes may be under selective pressure governed by parental-conflict (Haig and Westoby, 1989). In our study, we find a great overrepresentation of MEGs compared to PEGs in our reassessment of imprinting which suggest that MEGs may be under stronger selective pressure than PEGs according in support of the co-adaptation theory (Wolf and Hager, 2006). The data presented here further supports the differential dosage hypothesis (Dilkes and Comai, 2004), reflected by the limited occurrence of strict mono-allelic gene expression. Collectively these observations suggest that imprinting may be a product of several different selective pressures, and thus also allow genes of various functions to be imprinted.

## Materials and Methods

### Plant Material, Growth Conditions and Tissue Handling

All mutant lines used in this study are in the Col-0 background. Plants were grown on a 16 hrs light/ 8 hrs night cycle at 18 degrees. Accessions and mutants used; L*er*-1, Col-0, Tsu-1, *nrpE1* (SALK_029919) *drm1:drm2* (N16383) *mea-9* (SAIL_724_E07) *met1-3* (Saze et al., 2003), *met1-7* (SALK_076522), *rdr6-15* (SAIL_617_H07). For crosses, closed flower buds were emasculated 3 days prior to crossing to avoid self-pollination. The emasculated buds were left to mature. Upon crossing, pollen from the designated paternal donor was applied to the mature stigmas. For RNA extraction, siliques were dissected under a stereomicroscope and seeds were harvested into liquid nitrogen. Seeds from 3 plants, 4 siliques each were pooled in to each biological replicate. For phenotypic analysis of seed development, crosses and silique dissection was performed as described above. Seeds were mounted on a microscopy slide in a clearing solution of glycerol and chloral hydrate as previously described (Grini et al., 2002). Microscopy analyzes were performed using Zeiss Axioplan Imaging2 microscope system equipped with Nomarski optics.

### RNA Extraction, Probe Library Design and Preparation of Sequencing Libraries

Tissue was collected to MagNA-Lyser Green Beads Tubes (Roche) and RNA was extracted using SIGMA-ALDRICH Spectrum Plant Total RNA kit. Total RNA was quality checked by Nanodrop and Agilent RNA6000 kit for Agilent 2100 Bioanalyzer. For each sample, 100ng total RNA was spiked with ERCC Spike-In Control Mix (Ambion by Life Technologies) according to manufacturer’s protocol. The spiked total RNA was fragmented at 94 °C for 8 minutes to yield 100-200 bp fragments. cDNA libraries were prepared by using KAPA Stranded RNA-Seq Library Preparation Kit for Illumina platforms. Illumina TruSeq Adapters (SeqCap Adapter KitA (Roche)) were used to allow multiplexing. Probes between 50-105 bases (average length 80) were designed by Roche-Nimblegen based on TAIR10 sequences. Probe design failed for three genes: AT2G07739, AT3G19080 and AT5G58190. Final probe library is designed to capture 1011 genes, in total 5510 exons, probes were also designed to capture the RNA Spike-Ins. Probes were added to the multiplexed cDNA library and left to hybridize overnight at 47 °C. SeqCap Capture Beads (Roche) were used to recover hybridized cDNA. Agencourt AMPure XP Beads (Beckman Coulter) were used to purify the amplified cDNA libraries. 6-plexed cDNA libraries were sequenced using Illumina HiSeq 4000 System. Three biological replicates were sequenced for all crosses performed.

### Bioinformatics Analyzes

Illumina sequencing was used to analyze 54 samples representing three biological replicates each of 18 crosses (3 homozygous, 15 heterozygous); see SData 2. One Illumina sequencing library was prepared for each sample with a 300bp insert size target. Libraries were sequenced to produce 2x150 paired reads.

Reads were trimmed to remove low-quality bases and adapter sequences. Reads were first processed as pairs with cutadapt (Martin, 2011) version 1.8.1 using Illumina TruSeq adapter sequences. Reads were trimmed again with trimmomatic (Bolger et al., 2014) version 0.35 with parameters ILLUMINACLIP: TruSeq3-PE-2.fa:2:30:10 SLIDINGWINDOW:4:5 LEADING:5 TRAILING:5 MINLEN:100.

Reference coding sequences were selected for the 1011 genes under study. The TAIR10 version of *A. thaliana* Col-0 coding sequence (CDS) was downloaded in FASTA format from TAIR (Lamesch et al., 2012). A sequence subset was extracted to correspond to the 1011 genes with amplicons used in this study. The transcript with each [LOCUS].1 accession was used for each gene, with one exception: the transcript AT5G12170.2 was used because AT5G12170.1 had been deprecated in TAIR10. The result was a file of 1011 representative Col-0 transcript sequences (SGuideRawData 1).

The reads were initially mapped to the *A. thaliana* Col-0 reference CDS sequences. Read pairs were mapped such that at most one best map would be reported per read. The mapping used bowtie2 (Langmead and Salzberg, 2012) version 2.2.5 with parameters -p 4 --no-unal --no-mixed --no-discordant -q --phred33 -k 1 --end-to-end. The map process was launched on an SGE compute grid (SGuideProgramFiles 1). Counting pair-concordant maps only, the map rate was 55% for Col x Col pairs, 53% for L*er* x L*er* pairs, and 48% for Tsu x Tsu pairs; see SData 2. New ecotype specific consensus sequences were computed from the mappings of reads from the homozygous crosses. Each new reference sequence was computed with the consensus polisher Pilon (Walker et al., 2014), version 1.18 with options --fix all --changes. The polish process ran on each of three homozygous crosses separately, using reads from all three replicates per cross (SGuideProgramFiles Files 1, 2). The map+polish process was run in four iterations, at which point it had appeared to converge; a test 5^th^ iteration produced reported only trivial changes. The results were saved as FASTA (SGuideRawData 2).

Reads from all crosses were mapped to one pair of new reference sequences (SGuideRawData 3). Each pair was dictated by the cross; reads from a cross of Col and L*er* backgrounds were mapped to the Col and L*er* reference sequences, while reads from a cross of Col and Tsu backgrounds were mapped to Col and Tsu reference sequences. Reads from the Col x Col cross were mapped to the Col and L*er* reference sequences and, separately, to the Col and Tsu reference sequences. These mappings used bowtie2 as before with one parameter change to allow up to two targets per read: parameter -k 2 (SGuideProgramFiles 3).

Informative Reads were extracted from the map results. An Informative Read was required to have: mapped concordantly with its pair (i.e. proper orientation and approximate spacing for the read pair), mapped to the same gene in both strains, both mapQ scores ≥ 5, and either one strain’s alignment spanning InDels while the other is InDel-free, or both alignments InDel-free but one spanning fewer mismatches (i.e. SNPs) (SGuideProgramFiles 4).

Informative Reads from homozygous crosses were used to estimate noise and create the following noise filters. Three filters were applied to the gene sets. One filter excluded genes providing fewer than 50 Informative Reads per replicate. A second filter excluded genes providing fewer than 200 Informative Reads per cross. A third filter excluded genes providing less than 5-fold difference between Informative Reads preferring the true parent over the false parent (SGuideProgramFiles 5). Counts per gene are available in SData 3. To demonstrate minimal impact, the thresholds are plotted beside the score distributions in SFigure 2.

Informative Reads from heterozygous crosses were used to detect allelic expression bias. The scripted process (SGuideProgramFiles 6-9) compared 3 replicates of maternal and paternal Informative Read counts over the complete set of filtered genes. The maternal Informative Reads counts were halved to normalize for the 2:1 maternal-to-paternal expression bias expected in endosperm. The limma package (Ritchie et al., 2015) for R was used to fit a generalized linear model (lmFit()), normalize (makeContrasts(), contrasts.fit()), and generate significance levels (eBayes(), topTable()). A significance filter of P<=0.05 was used to detect significant levels of allelic expression bias.

Directional effects were tested in 15 pairs of reciprocal crosses. To allow comparison of the same gene in multiple samples, each Informative Read count was normalized by the total Informative Reads from all genes from that sample. Each gene test used 12 normalized counts representing maternal and paternal reads from three replicates each of two crosses. ANOVA with a generalized multiplicative model was applied to two factors: the pair of parents that were crossed, and the direction of the cross. A strong interaction of both factors (P<0.05) indicated a directional parental effect on the gene. The scripted process (SGuideProgramFiles 10-14) used R (Team, 2017) (https://www.R-project.org/) version 3.4.3 to generate tables (SGuideRawData 7-9).

Based on results of the ANOVA-based directional tests, gene subsets were formed using the method of Schon & Nodine (2017). Each subset (conservative, moderate stringency, low stringency, and unique to seed coat) was re-analyzed with the limma-based process to detect allelic expression bias (SGuideRawData 10-17).

Absolute expression analysis used the ERCC spike-in sequences (Ambion, Life Technologies). The analysis used total read counts (not Informative Read counts) per gene (not per parental allele). The analysis used RPK normalization *i.e.* reads per kilobase of transcript, rounded to nearest integer. A pseudocount of 1 was substituted for any normalized read count less than 1. Reads from all homozygous and heterozygous crosses were mapped to the new consensus sequences and, separately, to the ERCC reference sequences. Counts were ERCC-normalized using the DESeq2 R library (Love et al., 2014). The scripted process stored counts with DESeqDataSetFromMatrix and normalized using estimateSizeFactors() with the ERCC genes designated as controls. Thus, read counts from sample pairs were adjusted so as to maximize the similarity of each ERCC count from both samples. The ERCC-normalized counts were processed with our limma-based process to detect genes with differential expression between selected pairs of crosses. An on-line ERCC file repository is available under Supplementary Data Online.

The IRP assay was compared to SNP-based bioinformatics on a set of 12 transcripts selected to include putative MEGs, PEGs, and BEGs with above-threshold read counts in our data. Using pairs of new consensus sequences representing the Col and L*er* or Col and Tsu variants of these transcripts, SNPs were discovered using show-snps from the MUMmer package (Kurtz et al., 2005). The 21bp centered on each SNP was used to search reads from heterozygous crosses. The differential expression analysis pipeline was applied to the IRP and SNP-based read counts and the Pearson Product-Moment Correlation Coefficient was measured on the fold change per gene as detected using the IRP vs. SNP-based counts.

### Supplementary Data Online

Supplementary Data files and Supplementary Guide files are available online in the folder “Supplements” at [https://github.com/PaulGrini/Hornslien]. Guide files will point towards raw data and scripts. In the folder “ERCC _Normalize”, the Homozygous, L*er*, and Tsu directories hold the intermediate results for homozygous crosses, heterozygous crosses involving Col and L*er* strains, and heterozygous crosses involving Col and Tsu strains, respectively. The result files in the Normalized directory contain P-values for the differential expression between selected pairs of crosses, given three replicates per cross, for each gene.

All sequences generated in this study have been deposited in the European Nucleotide Archive (http://www.ebi.ac.uk/ena) with accession numbers (xxx-xxx).

## Acknowledgements

We thank Daniel Bouyer for fruitful discussions.

## Supplementary Figure legends

**Supplementary figure 1: Short outline of the experimental and bioinformatics pipeline.** 4 days after pollination, tissue was harvested from homozygous, heterozygous and mutant crosses for RNA isolation and preparation of cDNA libraries using the probe capture technology (Roche) and Illumina sequencing. Reads from homozygous crosses were used to create new reference transcriptomes for our targets, this allowed individual references for Col, L*er* and Tsu ecotypes. Reads from heterozygous and mutant crosses were all mapped to the new reference sequence to allow detection of parental specific gene expression.

**Supplementary figure 2: Thresholds for gene inclusion in this study.** Reads from homozygous crosses were mapped to consensus sequences from their true parent and a false parent. Genes were scored according to how well the true parent of origin could be detected from the read mapping. **Curves:** Gene scores for 1011 genes were sorted lowest to highest and plotted. Curve color indicates score: total reads/gene (green), fewest reads in a replicate (blue), and least fold difference in a replicate (red). Line type indicates cross: ColxCol reads mapped to Col and L*er* (dark color, solid line), L*er*xL*er* reads mapped to L*er* and Col (dark color, dotted line), ColxCol reads mapped to Col and Tsu (light color, solid line), and TsuxTsu reads mapped to Tsu and Col (light color, dotted line). **Straight solid lines:** Minimum thresholds were selected near the first elbow of each curve. The thresholds are: minimum 200 reads/gene (green), minimum 50 reads/replicate (blue), and minimum 5-fold specific preference to true parent (red). Any gene with at least one score below any thresholds was excluded from further study. Note the y-axis uses a log scale and scores below 1 are not plotted.

**Supplementary figure 3: SNP vs. Informative Reads Pipeline (IRP).** The pipeline developed in this study was compared to the traditional method of counting reads mapping to a certain known SNP. Well established MEGs and PEGs were selected for this comparison as well as novel identified in this study and some biparentally expressed genes (BEGs) as negative controls. Reads mapping to a single SNP was chosen and processed in the same manner as the IRP. Correlation was high, although some outliers were observed, e.g. FIS2 and SKS14. The Pearson Product-Moment correlation coefficients (cor) for SNP versus IRP are displayed for each cross.

**Supplementary figure 4: Fold Change (FC) Distribution in the difference stringency groups. A)** FC-distribution in our full data set based on genes with informative reads and good mapping abilities show that many genes produce extreme FC-values. **B)** Removing seed coat contaminants from further analysis do indeed remove a big portion of the more extreme FC-values **C)** Further removal of possible seed coat contaminants due to no expression information in public data base gives a further reduction of extreme FC-values.

**Supplementary figure 5: Parentally biased expression in the conservative and low stringency group. A)** and **B)** The LogFC between maternal and paternal reads for each gene in each cross was plotted against its reciprocal counterpart to display preferential mapping, i.e. biased expression. A positive LogFC signifies maternal preference while a negative LogFC signifies paternal preference. **A)** The conservative stringency group and **B)** the low stringency group of targets for the two cross pairs; Col-L*er* (left panels) and Col-Tsu (right panels). Grey filled circles signify targets that do not have significant parentally biased expression. Red and blue filled circles have maternally or paternally biased expression, respectively, in both directions of a cross pair. Orange filled circles have parentally biased expression in only one direction of a cross pair. Black circles show opposite preferential expression depending on the direction of the cross. Cross direction is always indicated as ♀ x ♂ **C)** and **D)** Great overlap is observed between preferentially expressed genes detected in all ecotype crosses used in this study; Col-L*er* (C-L) reciprocal crosses and Col-Tsu (C-T) reciprocal crosses, Maternally Expressed Genes (MEGs) in left panels and Paternally Expressed Genes (PEGs) in right panels. **C)** 18 genes show the same preferential bias in all ecotypes studied in the conservative endosperm specific group of targets. **D)** 295 genes show parental bias in our low stringency group. Genes that do not overlap are most frequently genes that do not have sufficient SNP/InDels in the alternative cross pair and could not be assessed. Few genes show inconsistent or not reciprocal preferential expression in the two cross pairs used in this study. Grey circle; genes that show reciprocal parent preferential expression in Col-Tsu cross pair. Coloured peanut; genes that show reciprocal parent parental expression in Col-L*er* cross pair.

**Supplementary figure 6: No overlap between imprinted genes at globular versus torpedo stage in the conservative stringency set.** Comparing the conservative stringency set obtained by using same filtering conditions as a recent report (Schon and Nodine, 2017) we did not detect any overlap for maternally expressed genes (MEGs) or paternally expressed genes (PEGs).

**Supplementary figure 7: All crosses with mutant parents have reached the same developmental stage at four days after pollination. A)** Seeds from crosses between wild type and mutant lines used in this study were investigated at four days after pollination (DAP) indicating that there is no obvious difference regarding seed development depending on the specific mutants used as a maternal or paternal contributor in a cross. **B)** In around 80% of all seeds investigated, the embryos were classified as 32-cell to globular stage at 4DAP, and only very few deviated from this stage. Data is presented as means ±standard deviation across three biological replicates. Cross direction is always indicated as ♀x ♂

**Supplementary figure 8: Parental specific expression is changed in L*er*x*met1* and *mea-9*xL*er* crosses compared to wild type (WT). A)** LogFC ratios obtained from relative expression analysis of informative reads obtained from parentally biased genes in L*er*xCol WT compared to L*er*x*met1-7* show deregulation in mutant cross mainly for maternally expressed genes (MEGs). **B)** Analysis of LogFC ratios from parentally biased genes in ColxL*er* WT and *mea-9*xL*er* mainly show deregulation of paternally expressed genes (PEGs). Each locus investigated is represented by two circles oriented on the same vertical line. Purple and green circles represent LogFC ratios in mutant crosses; red and blue represent LogFC ratios in WT crosses.

**Supplementary figure 9: Mutants in RdDM are not sufficient to deregulate imprinted gene expression of paternally expressed genes (PEGs).** Ratios of the change of maternal reads in mutant (mat2): maternal reads in wild type (mat1) were used to create a prediction interval (purple square) for analysis of paternal ratios; paternal reads in mutant (pat2): paternal reads in wild type (pat1). Based on the prediction interval made on maternal reads, genes where maternal reads fell outside the interval were not considered as significant (red circles). Low degree of deregulation is observed of the paternal allele for PEGs in crosses with mutants of the RdDM pathway **A)** L*er*x*drm1;drm2* **B)** L*er*x*nrpe1*, **C)** L*er*x*rdr6*. **D)** Limited overlap of the few deregulated genes in the RdDM mutant crosses of **A)**, **B)** and **C)**.

**Supplementary figure 10: Limited deregulation of maternally expressed genes (MEGs) in crosses with maternal mutants of RdDM.** Ratios of the change of paternal reads in mutant (pat2): paternal reads in wild type (pat1) were used to create a prediction interval (purple square) for analysis of maternal ratios; maternal reads in mutant (mat2): maternal reads in wild type (mat1). Most genes cluster inside the prediction interval and a low degree of deregulation is observed in all crosses **A)** *drm1;drm2*xL*er*, **B)** *nrpe1*xL*er*, **C)** *rdr6*xL*er*. Based on the prediction interval made on paternal reads, genes where paternal reads fell outside the interval were not considered as significant; blue circles in **A)**, **B)** and **C)**. **D)** The intersection of genes where the maternal allele is downregulated is restricted to one gene (AT4G00780).

**Supplementary figure 11: Limited deregulation of paternally expressed genes (PEGs) in crosses with maternal mutants of RdDM.** Ratios of the change of paternal reads in mutant (pat2): paternal reads in wild type (pat1) were used to create a prediction interval (purple square) for analysis of maternal ratios; maternal reads in mutant (mat2): maternal reads in wild type (mat1). Most genes cluster inside the prediction interval and a low degree of deregulation is observed in all crosses **A)** *drm1;drm2*xL*er*, **B)** *nrpe1*xL*er*, **C)** *rdr6*xL*er*. Based on the prediction interval made on paternal reads, genes where paternal reads fell outside the interval were not considered as significant; blue circles in **A)**, **B)** and **C)**. **D)** The intersection of genes where the maternal allele is upregulated shows no overlap between crosses.

**Supplementary table 1: Embryo development statistics in wild type crosses.** Siliques were dissected, and seeds were analysed in a regular light microscope to score embryo development at 4 days after pollination as well as number of fertilized seeds. Embryos were classified from 4-celled or less through transition/early heart. Data is presented as means ± standard deviation (SD) across three biological replicates, n depicts total number of seeds analysed. Cross direction is always indicated as 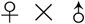.

**Supplementary table 2: Genes displaying opposite preferential expression. A) Col-L*er* reciprocal crosses and B) Col-Tsu reciprocal crosses.** Differential expression (DE) fold change (FC) values are given for maternal-paternal preference. Positive FC value signifies maternal preference, negative FC signifies paternal preference. Homozygous crosses were compared in a separate DE analysis based on listed average normalized read counts giving the ERCC FC in L*er* or Tsu compared to Col.

**Supplementary table 3: Possible candidates for ecotype specific imprinting. A) L*er*xCol and TsuxCol B) ColxL*er* and ColxTsu.** Overlapping genes that display non-reciprocal preferential expression comparing the two cross pairs Col-L*er* and Col-Tsu. Positive FC value signifies maternal preference, negative FC signifies paternal preference. Genes depicted in bold show preferential mapping dependent on ecotype. Imprinting state reported in selected whole genome surveys of imprinting: H= Hsieh et al., 2011; G= Gehring et al., 2011; W= Wolff et al., 2011; P= Pignatta et al., 2014; S= Shirzadi et al., 2011 downregulated (x) in *cdka;1*-screen.

**Supplementary table 4: Genes showing maternal and paternal preferential expression in the conservative stringency set. A) Maternally expressed genes (MEGs), B) Paternally expressed genes (PEGs).** Differential expression (DE) fold change (FC) values are given for maternal-paternal preference for ColxL*er*, L*er*xCol, ColxTsu and TsuxCol crosses. Positive FC value signifies maternal preference, negative FC signifies paternal preference; P-values obtained from DE analysis ranged from 1,18x10^-94^ to 0,0059 (MEGs) and 1,55x10^-101^ to 0,0001 (PEGs). Imprinting state reported in selected whole genome surveys of imprinting (x if consistent with this data): H= Hsieh et al., 2011; G= Gehring et al., 2011; W= Wolff et al., 2011; P= Pignatta et al., 2014; S= Shirzadi et al., 2011 downregulated (d) in *cdka;1*-screen. CLT= SNPs in all ecotypes tested, CL= only SNP between Col-0 and Ler, CT= Only SNP between Col-0 and Tsu.

**Supplementary table 5: Paternal preferential expression in the moderate stringency set.** Differential expression (DE) fold change (FC) values are given for maternal-paternal preference for ColxLer, L*er*xCol, ColxTsu and TsuxCol crosses. Negative FC signifies paternal preference. P-value range obtained from DE analysis ranged from 1,15x10^-7^ to 0,0099. Imprinting state reported in selected whole genome surveys of imprinting (x if consistent with this data): H= Hsieh et al., 2011; G= Gehring et al., 2011; W= Wolff et al., 2011; P= Pignatta et al., 2014; S= Shirzadi et al., 2011 downregulated (d) in *cdka;1*-screen. Expression pattern indicated is based on Microarray ATH1-chip library (Belmonte et al., 2013) and re-analysis (see Material and Methods) for tissue enrichment to identify endosperm specific (ess.) expression compared to all other seed compartments at globular stage, unless otherwise indicated (PG=Preglobular, LC=Linear Cotyledon, BC=Bent Cotyledon). ND=not detected in seed. CLT= SNPs in all ecotypes tested, CL= only SNP between Col-0 and L*er*, CT= Only SNP between Col-0 and Tsu.

**Supplementary table 6: Maternal preferential expression in the moderate stringency set.** Differential expression (DE) fold change (FC) values are given for maternal-paternal preference for ColxL*er*, L*er*xCol, ColxTsu and TsuxCol crosses. Positive FC value signifies maternal preference, P-values from DE analysis were all <0,01. Imprinting state reported in selected whole genome surveys of imprinting (x if consistent with this data): H= Hsieh et al., 2011; G= Gehring et al., 2011; W= Wolff et al., 2011; P= Pignatta et al., 2014; S= Shirzadi et al., 2011 downregulated (d) in *cdka;1*- screen. CLT= SNPs in all ecotypes tested, CL= only SNP between Col-0 and Ler, CT= Only SNP between Col-0 and Tsu.

**Supplementary table 7: Embryo development statistics in mutant crosses.** Siliques were dissected, and seeds were analysed in a regular light microscope to score embryo development at 4 days after pollination as well as number of fertilized seeds. Embryos were classified from 4-celled or less through transition/early heart. Data is presented as means ± standard deviation (SD) across three biological replicates, n depicts total number of seeds analysed. Cross direction is always indicated as 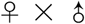.

**Supplementary table 8: Paternal read change of maternally expressed genes (MEGs) in crosses to *met1*.** Average normalized paternal read counts across three biological replicates, in L*er*xCol and TsuxCol compared to the respective met1 mutant cross, and the associated ratios. For all read counts reported, a students t-test showed significant change in mutant compared to WT (p-values <0,05). Some genes fail the prediction interval test (indicated by fail tests) because the maternal reads are outside the initial prediction interval, paternal reads from these were not analysed. CLT= SNPs in all ecotypes tested, CL= only SNP between Col-0 and L*er*, CT= Only SNP between Col-0 and Tsu.

**Supplementary table 9: Imprinted genes not affected by mutation in *MET1* and *MEA*. A) MEGs not regulated by MET1 in pollen, B) PEGs not affected by mutation of MEA in seed parent.** Differential expression (DE) fold change (FC) values are given for maternal-paternal preference as an average of wild type crosses ColxL*er*, L*er*xCol, ColxTsu and TsuxCol. Imprinting state reported in selected whole genome surveys of imprinting (x if consistent with this data): H= Hsieh et al., 2011; G= Gehring et al., 2011; W= Wolff et al., 2011; P= Pignatta et al., 2014; S= Shirzadi et al., 2011 downregulated (d) in cdka;1-screen. CLT= SNPs in all ecotypes tested, CL= only SNP between Col-0 and Ler, CT= Only SNP between Col-0 and Tsu. *Not analyzed, maternal reads outside prediction interval; **Paternal reads downregulated; ***Maternal reads downregulated.

**Supplementary table 10: Dynamic regulation of paternally expressed gens by MET1, PRC2 and possibly RdDM.** x=deregulated in mutant cross. CLT= SNPs in all ecotypes tested, CL= only SNP between Col-0 and L*er*, CT= Only SNP between Col-0 and Tsu.

